# Gene clustering drives the transcriptional coherence of disparate biological processes in eukaryotes

**DOI:** 10.1101/2021.04.17.440292

**Authors:** Richard I. Joh, Michael S. Lawrence, Martin J. Aryee, Mo Motamedi

## Abstract

The establishment of distinct transcriptional states entails coordinated transcriptional coregulation of several disparate biological processes (BP), involving thousands of genes scattered throughout the genome. Linear clustering of genes in a single BP is one strategy for their transcriptional coregulation. However, whether such gene clustering also plays a role in transcriptional coherence of several disparate BPs remains unexplored. Here, by analyzing the genomes of eukaryotes ranging from yeast to human, we report the identification of thousands of conserved and species-specific disparate clustered BP pairs, many of which in normal human tissues are transcriptionally correlated. Strikingly, our results reveal that often this system-level transcriptional coordination is achieved in part by the genic proximity of regulatory nodes of disparate BPs whose coregulation drives the transcriptional coherence of their respective pathways. This, we hypothesize, is one strategy for creating coregulated, tunable modulons in eukaryotes.

## Introduction

To survive stress or deploy developmental programs, cells establish a spectrum of distinct transcriptional states. This involves the rapid, robust and reproducible transcriptional coordination of hundreds or thousands of genes dispersed throughout the genome governing dozens of disparate biological processes (BPs). To study this type of system-level change in biological outputs, gene regulatory networks (GRNs) were developed to functionally and temporally link the regulatory genes and signaling components of a multitude of disparate BPs^1–6^. These networks are modular and hierarchical^7,8^, within which several overrepresented network motifs or sub-circuits^9,10^, create positive and/or negative regulatory loops through which biological outputs of up to thousands of genes are coordinated. Redundancies, stratifications and amplifications of motifs within GRNs not only allow the incorporation of signals from a variety of cellular and environmental stimuli^7^, but also together produce robust, timely, fine-tuned and stable biological responses. Indeed, the system-level coregulation achieved within GRNs plays a central role in adaptation and development and thus is under strong evolutionary selection^11–13^. Organizationally, GRNs are made up of lower-level sub-circuits, such as regulons - a set of genes that are transcriptionally coregulated as a unit - and higher-level sub-circuits, such as modulons, often comprised of several regulons that become transcriptionally linked in response to the same stimuli. Mechanistically, the lower level sub-circuits are formed by the combinatorial activities of *trans*-acting factors, such as transcription factors (TF), and chromatin regulatory proteins targeted to a given set of *cis*-regulatory DNA sequences^12,14^.

Gene clustering is one means by which the transcription of a pair or a large group of interrelated genes can be coordinated. This was first described in prokaryotes where functionally interrelated genes cluster into operons from which poly-cistronic messenger RNAs (mRNAs) are transcribed, ensuring their co-expression^15^. Even though poly- or di- cistronic mRNAs of protein coding genes are rarely found in metazoans except for nematodes^16–18^ and flies^19–21^, genomic clustering of interrelated genes is a prevalent and conserved feature of eukaryotic genomes in organisms such as yeast^22–24^, fly^25–27^, worm^28–30^, zebrafish^31,32^, mouse^33,34^ and human^34–41^. In fact, eukaryotic genomes possess multiple domains of transcriptional coregulation^25,27,40^, within which a shared chromatin state and *cis*-regulatory elements help establish the transcriptional activities of these regions^42–44^. This type of gene clustering has functional consequences. For example, gene clustering helps the transcriptional coordination (e.g. housekeeping and highly expressed genes) and temporal regulation (e.g. HOX family of TFs) of several groups of interrelated genes^27,35,45–48^. Clearly, this type of proximity-driven transcriptional co-regulation plays an important role in life, evidence by the fact that its disruption is linked to several developmental pathologies in human^49–52^.

Beyond the linear placement of genes in the genome, distant genes can be found in close contact to other co-regulated genes, forming larger co-localized clusters in the nuclear three dimensional (3D) space^53–58^. This helps coordinate the transcription of genes at both the micro- (neighboring genes) and macro-scales (A/B compartments and topologically associated domains (TADs))^59–65^. Similarly, their physical proximity in the nucleus helps establish a common chromatin state and enable a common transcriptional output among interrelated genes^66^. Together these discoveries demonstrate that genome organization is not random and that clustering of interrelated genes is a highly conserved strategy for driving their coregulation. However, whether the linear placement of unrelated genes helps couple the transcriptional output of disparate GRN sub-circuits remain an area of active investigation.

Previously, using the fission yeast (*Schizosaccharomyces pombe*), we showed that as cells enter quiescence (G0), the constitutive heterochromatin protein Clr4 - the sole lysine 9 histone H3 (H3K9) methyltransferase in this organism - is deployed to euchromatic parts of the genome to coregulate the expression of hundreds of genes^67^. Unexpectedly, we found that most of these coregulated genes occur in linear gene arrays, and were together overrepresented in disparate BPs such as development, cell cycle and metabolism, whose repression is important for establishing the G0 state^68^. In addition to examples in yeast^69,70^, formation of transcriptionally coregulated linear gene arrays are also found in mammals. For example, growth factor-induced exit from quiescence of mouse fibroblast cells also results in the time-dependent transcriptional coregulation of linear gene arrays^42^. These and other observations^71,72^ together with our work in the fission yeast suggested that clustering of genes belonging to disparate BPs within a few gene neighborhoods may be a conserved strategy for efficiently targeting the same transcriptional regulatory proteins through which linear sections of the genome can become coregulated. If true, then this type of genome architecture would couple the transcription of disparate lower level GRN sub-circuits, help create modulons and would thus be under strong evolutionary selection.

Here to test this hypothesis, we devised a statistical framework to quantify genomic clustering between disparate BPs in five of the most commonly used and best-annotated eukaryotic model organisms ranging from yeast to human spanning over one billion years of evolution^73,74^. Using this platform, we identified thousands of conserved and species-specific clustered BP pairs, many of which are strongly transcriptionally correlated in normal human cells. Interestingly, even in the absence of pathway-wide clustering, disparate BP pairs sharing at least one clustered TF pair display strong transcriptional coherence, suggesting that clustered TFs represent nodes of transcriptional coregulation, through which transcriptional output of their associated BPs can be coupled. In support of this, we find that disruption to TF clustering (deletion or translocation) results in the loss of the transcriptional coherence of associated BPs. Overall, these data suggest that genic proximity of regulatory nodes of disparate BPs can couple their transcription, thus enabling the establishment of tunable, coregulated modulons in eukaryotes.

## Results

### Identification of clustered BP pairs in eukaryotic model organisms

Our work in the fission yeast led us to hypothesize that in eukaryotes, clustering of genes belonging to disparate BPs may help their transcriptional coregulation. This could help create important modulons and would be thus under selection. To identify such BP pairs (Figure 1A), we developed a statistical framework to quantify clustering based on the positioning of all protein-coding genes (Figure 1B) in the fission yeast(*Schizosaccharomyces pombe*), nematode (*Caenorhabditis elegans*), fruit fly (*Drosophila melanogaster*), mouse (*Mus musculus*) and human (*Homo sapiens*) genomes. The highly annotated genomes of these organisms endowed our analyses with the power to examine thousands of conserved and species-specific biological pathways across evolution (Figure 1C). BP terms in the Gene Ontology (GO) consortium^75^ were used as a means to define biological pathways in our analyses. Uniquely, GO provides an up-to-date, pan-organismal classification of BP terms in the aforementioned organisms, which are comparable across species. (We also used other biological pathway classification platforms, including KEGG, BioCarta, Wiki Pathways, to confirm our findings in the human genome (See Figures S12-S13)).

**Figure 1.**
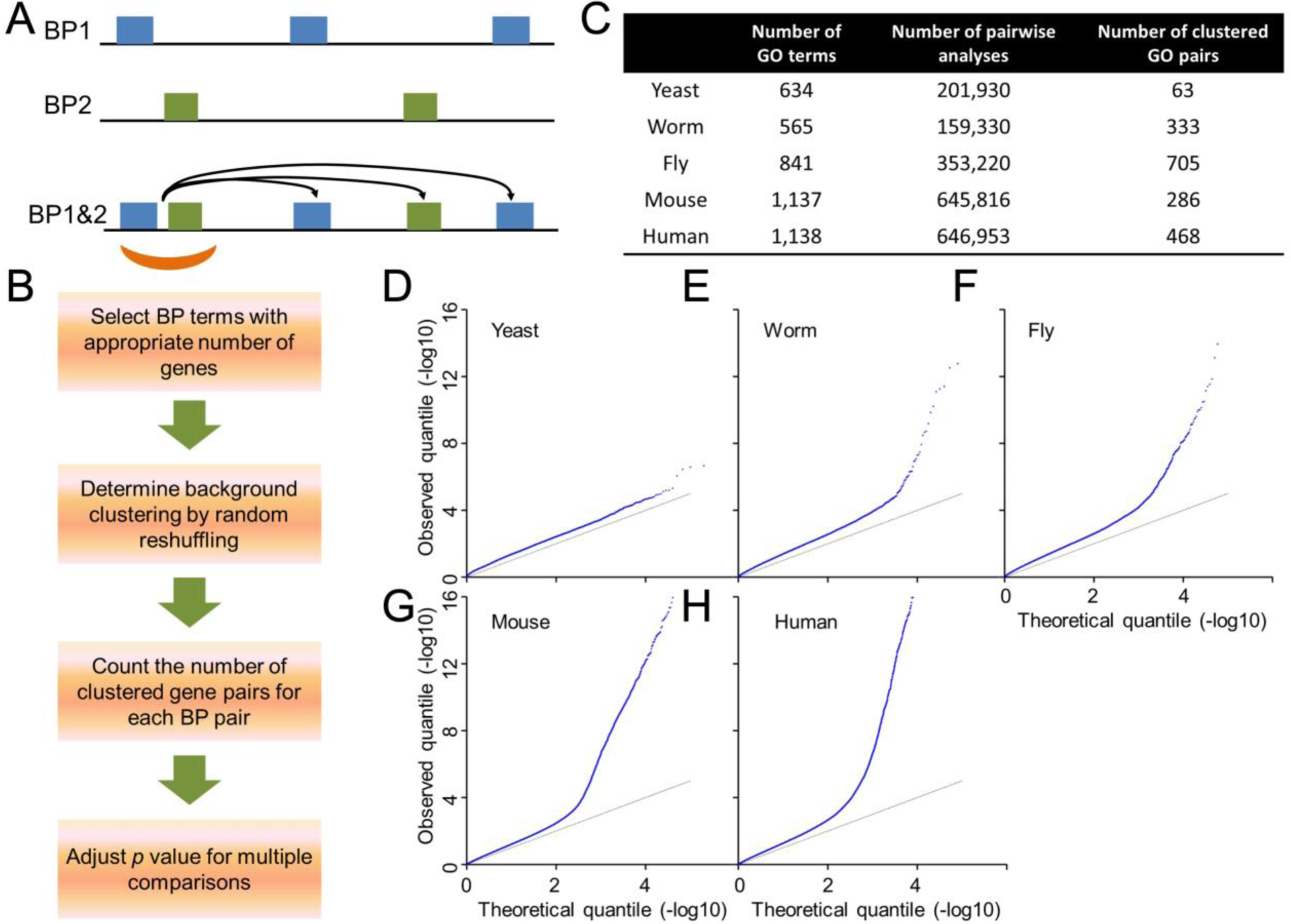
Hundreds of disparate Gene Ontology (GO) biological process (BP) pairs are clustered in the genomes of organisms ranging from yeast to human. (A) Scheme depicting the hypothetical distribution of two BPs and their associated genes on a chromosome. Blue and green boxes depict genes in BP1 and BP2, respectively. The orange arc denotes a clustered gene pair. The arrows indicate the direction of transcriptional regulation assuming that the clustered genes are both transcription factors. (B) Flow chart of all analytical steps in this study. (C) Table summarizing the numbers of BP terms, pairwise analyses, and significantly clustered BP pairs identified in each organism. (D-H) Quantile-quantile (Q-Q) plots of observed (Y-axis) versus theoretical (X-axis) of *p*-value distributions for BP-BP clustering in (D) yeast (*S. pombe*), (E) worm (*C. elegans*), (F) fly (*D. melanogaster*), (G) mouse (*M. musculus*) and (H) human (*H. sapiens*).

For the detailed description of the statistical method developed for identifying significantly clustered BP pairs, please see (Figure 1A-C) and associated sections in StarMethods. Briefly, we (1) selected BP terms with sufficient number of genes to provide enough statistical power for our analyses in each genome (Table S1), (2) set a threshold distance for gene clustering that captures the majority of the proximity-driving gene-pair transcriptional correlations (Figure S1 and StarMethods), and tested its statistical robustness against a range of alternative threshold distances (Figure S2-3), (3) demonstrated the specificity (Figure S4) and performance (Figure S5) of the statistical method using randomly generated gene sets (see StarMethods), (4) determined background clustering by performing 2,000 random samplings (Figure S6) for each BP- BP analysis in each organism, and (5) estimated *p* values for each BP-BP analysis in each organism (Table S2). The *p*-values were used to generate quantile-quantile (QQ) plots shown in Figure 1D-H. The QQ plot for each organism stayed close to the expected diagonal, indicating minimal systematic inflation or deflation. However, the tails of the QQ plots curl upwards in all organisms except the fission yeast indicating that metazoan genomes have thousands of significantly clustered BP pairs. The peculiarity of the fission yeast QQ plot is likely the result of its distinct genome organization relative to metazoans: (A) gene density of protein-coding genes is almost uniform across the fission yeast euchromatic domains, and (B) 50% of all fission yeast protein-coding genes overlap with another gene. Overall, these plots reveal that the statistical framework developed here for capturing BP-BP clustering works effectively in metazoans but may not be suited ideally for the uniformly dense and highly overlapped genomes of yeast species. Next, for each organism, we adjusted the significance of clustering for multiple comparisons using positive false discovery rate^76^. In the end, out of over two million BP-BP analyses, 1,855 BP pairs were found to be significantly clustered (*FDR* < 0.05) in the five genomes analyzed (Figure 1C, Table S2). Figure 2 shows examples of clustered GO pairs in each organism. Even though some of the clustered BP pairs identified are functionally similar (fs), many BP pairs are functionally disparate (fd) (Table S2). Together, these analyses demonstrate that genome clustering of fd-BPs is an organizational feature of eukaryotic genomes.

**Figure 2.**
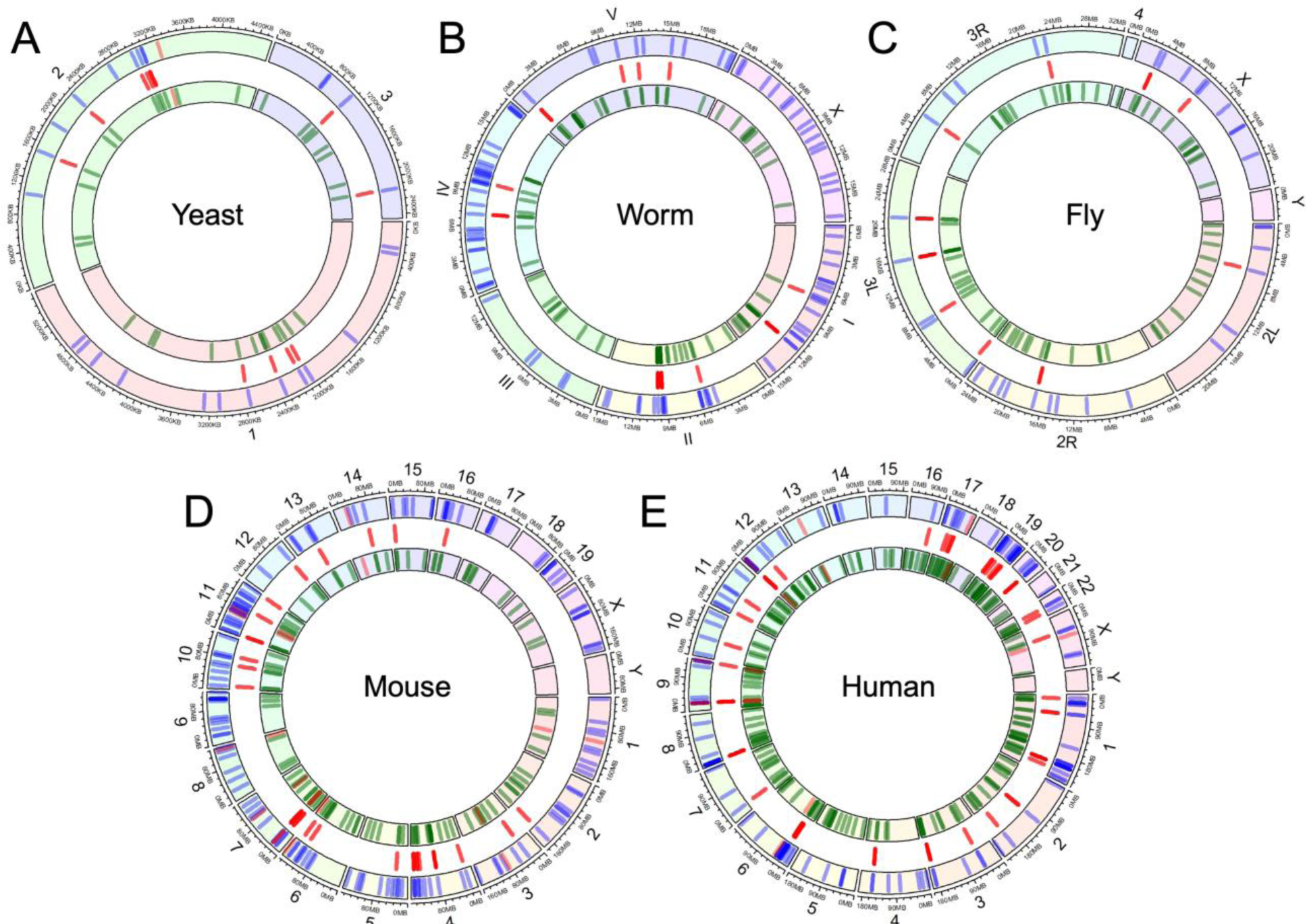
Representative clustered BPs in organisms ranging from yeast to human. Ring plots depicting representative clustered BP pairs in (A) yeast (*S. pombe* GO:0032956 (regulation of actin cytoskeleton organization) and GO:2001251 (negative regulation of chromosome organization), (B) worm (*C. elegans* GO:0007169 (transmembrane receptor protein tyrosine kinase signaling pathway) and GO:1901136 (carbohydrate derivative catabolic process)), (C) fly (*D. melanogaster* GO:0045747 (positive regulation of Notch signaling pathway)and GO:0046486 (glycerolipid metabolic process)), (D) mouse (*M. musculus* GO:0030308 (negative regulation of cell growth) and GO:0033135 (regulation of peptidyl-serine phosphorylation)) and (E) human (*H. sapiens* GO:0006959 (humoral immune response) and GO:0007162 (negative regulation of cell adhesion)). Ring segments with a number on the outside denote a different chromosome. Blue and green lines in outer and inner rings indicate the genomic location of each gene in their respective pathways. Red lines in central gap indicate locations of clustered gene pairs.

### Several similar BP pairs show conserved clustering in eukaryotic genomes

If clustering of genes belonging to fd-BPs couples their transcription and helps create GRNs, then its maintenance would be under selection in evolution. Consistent with this hypothesis, 17 identical significantly clustered BP pairs were identified in the mouse and human genomes (Table S2). This is largely driven by synteny between these two species which shared a recent common ancestor^77^. Interestingly, we also found that several clustered BP pairs, though not identical, were highly similar to one another across these genomes, suggesting that such BP clustering is under selection and thus maintained in evolution.

To determine whether similar pairs of BPs cluster in the genomes of these distantly related eukaryotes, we selected BPs that have a clustered BP partner in at least two organisms (See StarMethods). The cutoff of two permits the identification of mammalian-specific (mouse and human) or metazoan-specific (fly and worm) clustered BP pairs. This yielded 393 such BP pairs (Table S2, and StarMethods). We then applied semantic similarity^78^ developed by Lin^79^ or Resnik^80^ and grouped the 393 BP pairs into 30 highly similar BP groups (Table S3 and Figure 3A). Semantic similarity quantifies (range 0.0 - 1.0) the functional distance between two BP terms based on their nested organization in the GO tree, depth and context relative to one another. The two semantic similarity methods produced highly overlapping BP groupings (Figure S7A), suggesting their robustness in organizing these BP terms into similar BP groups. We then asked in how many different genomes BPs in each group cluster with BPs in other groups. This analysis revealed several highly similar clustered BP pairs in the five eukaryotic genomes (Figure 3B and Figure S8).

**Figure 3.**
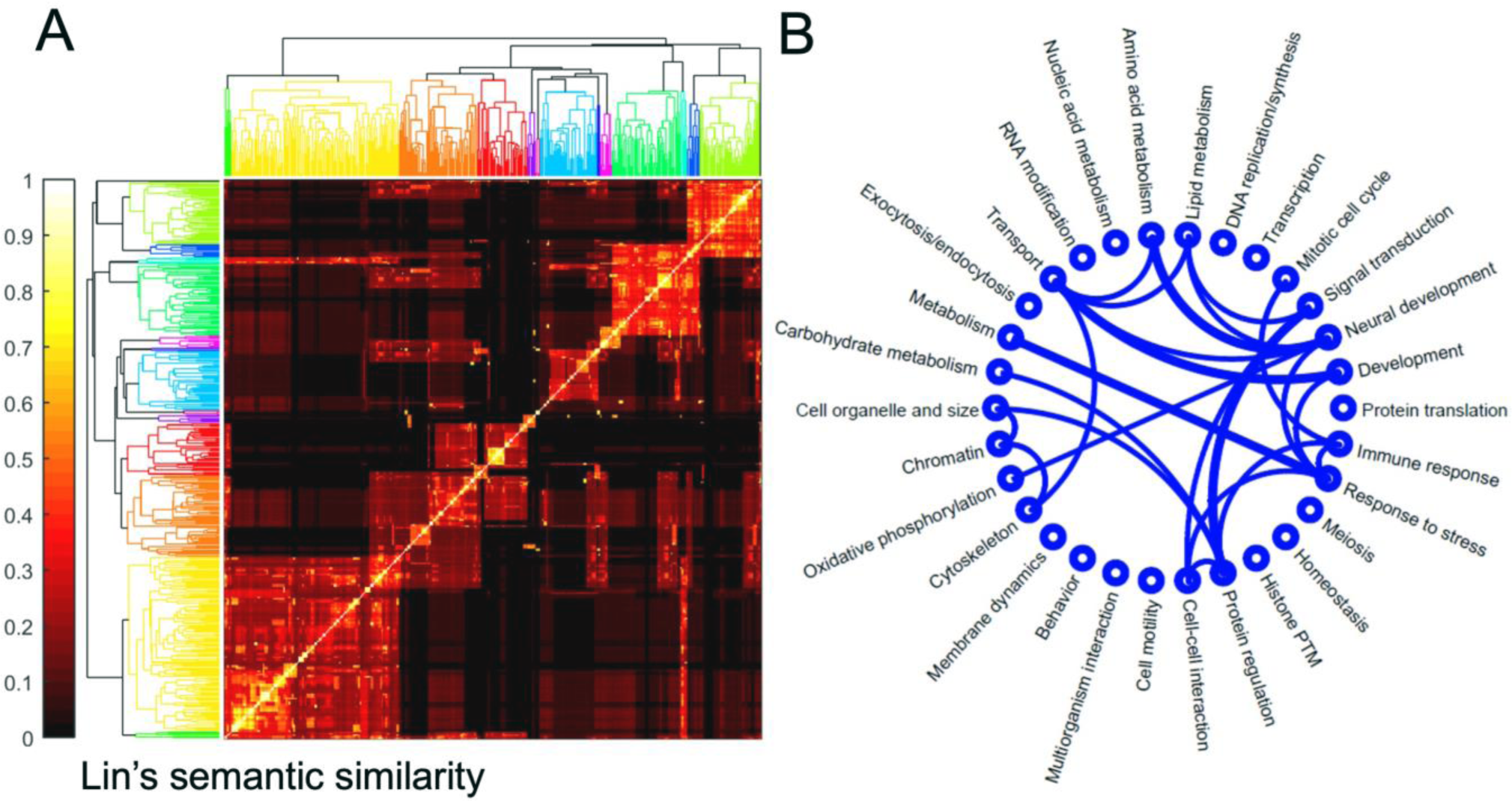
Several similar BP pairs show conserved clustering in eukaryotes. (A) Heatmap depicting hierarchical clustering of BP terms with clustered BP partners in at least two model organisms. Color represents the Lin’s semantic similarity score (0 - 1.0). (B) Visual representation of clustering among different BP groups in yeast, worm, fly, mouse, and human. Lines connecting two BP groups indicate clustering in at least three genomes. Thickness of lines is proportional to the number of organisms (three or more) in which pairwise clustering occurs.

To further test this hypothesis in parallel, we also asked whether a given BP in Table S3 clusters with similar BP terms in the other genomes. For example, if BP1 clusters with BP2 and BP2’ in mouse and yeast genomes, respectively, we calculated the BP2-BP2’ similarity by Lin. Our results (Figure S7B) revealed that the frequency with which highly similar (Lin scores >0.9) BP2-BP2’ are found is significantly higher than background, supporting the hypothesis that a given BP tends to cluster with highly similar BPs in multiple eukaryotic genomes. For example, metabolism BPs cluster with stress response BPs in four out of the five genomes, consistent with the well-established link between these two processes in biology^81–83^. Another example is the conservation of clustering between the amino acid metabolism and neural development BPs. Considering the critical role that amino acids and their metabolites play in synaptic transmission^84,85^, and learning and memory^86,87^, our data suggest that this type of genomic clustering may portend a functional link in their coregulation in organisms ranging from fly to human. We also noted several conserved clustered BP pairs which currently do not have an obvious functional link (Table S3). These suggest hitherto unknown functional relationships between these BP pairs (see Discussion for additional examples). Overall, these analyses demonstrate that clustering of hundreds of BPs is maintained over long evolutionary timescales, suggesting that this type of genome organization may enable their transcriptional co-regulation.

### Clustered BPs are transcriptionally correlated

The hierarchical or nested organization of GO contains many functionally similar (fs) or overlapping BP terms which are predicted to be transcriptionally coupled. We, on the other hand, would like to determine whether genome clustering could couple the transcriptional output of functionally disparate (fd) or non-overlapping BP pairs. To perform this analysis, we first developed a pipeline for quantifying BP-BP transcriptional correlation by calculating Z scores for all genes across the 53 different tissues represented in the GTEx dataset^88^ (Figure S9A-D). BP-BP correlations were calculated after removing shared genes and clustered gene pairs whose inclusion would artificially increase the transcriptional correlation between two BPs (Figure S1). Next, we used semantic similarity to define fd-BP pairs as those with low semantic similarity scores (Lin score ≤0.1). As expected, fs-BP pairs display higher transcriptional correlation relative to fd-BP pairs (Figure S10A-B). Moreover, comparing the transcriptional correlations of fd- BP pairs revealed that clustered fd-BP pairs (N=278) display higher transcriptional correlation compared to unclustered fd-BP pairs (N=458,999) (see StarMethods), supporting a model in which clustering of fd-BPs links their transcription. Also, because shared genes and clustered gene pairs were removed from our correlation analyses, we conclude that the correlations among clustered fd-BP pairs are pathway-wide, and not driven the coincidental correlation of clustered gene-pairs.

### BP pairs which share a clustered TF display significant transcriptional correlation

Interestingly, several of the clustered gene pairs in clustered fd-BPs were TFs. Because regulatory nodes of GRNs often consist of *trans*-acting co-expressed genes^89,90^, this prompted us to test whether clustered TFs are a strong predictor of the transcriptional coherence of clustered BP pairs. We divided clustered fd-BP pairs (N=278) into three groups: 1) one or both BPs lacks an annotated TF (N=18); 2) both BPs contain at least one annotated TF, and the TFs are not clustered (N=123); 3) both BPs contain at least one annotated TF, and at least one pair of TFs is clustered (N=142). Our analysis revealed that clustered fd-BPs with clustered TFs display the highest transcriptional correlation of the three groups (Figure 4B). Figure 4C shows an example of a clustered BP pair with clustered TFs in human, namely humoral immune response (GO:0006959) and negative regulation of cell adhesion (GO:0007162) (also shown in Figure 2E). Likewise, clustered fs-BP pairs (Lin score>0.1) displayed higher transcriptional correlation relative to unclustered fs-BP pairs (Figure S10C), especially those with a clustered TF pair (Figure S10D). Together, these data support a model in which clustered TFs can be used to couple the transcription of their associated BPs in human.

**Figure 4.**
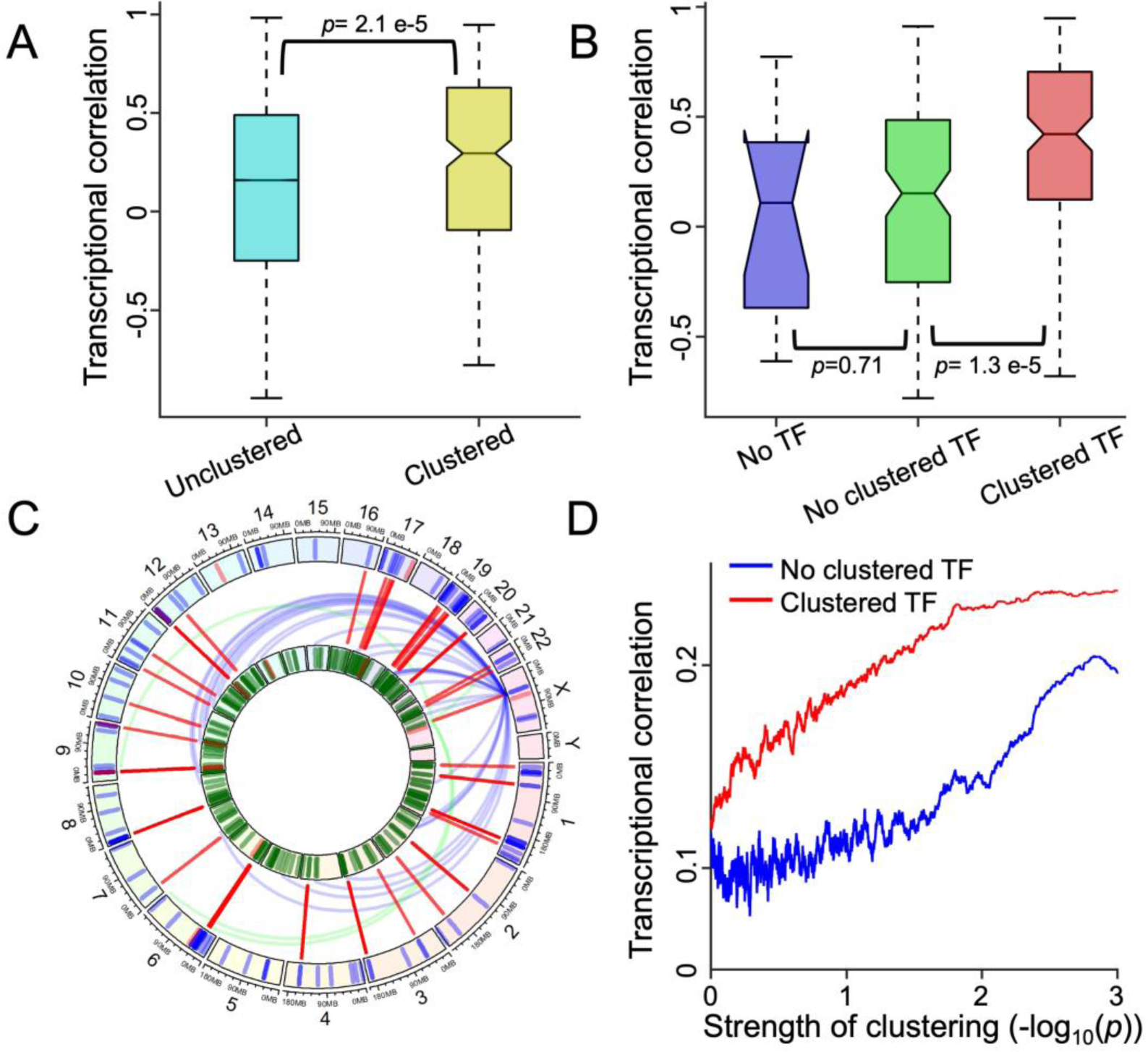
Functionally disparate (fd) clustered BP pairs (Lin Score. ≤**0.1), especially those with clustered TFs, are transcriptionally correlated in human.** (A) Box plots depicting transcriptional correlation of clustered (N=458,999) and unclustered (N=278) fd-BP pairs. (B) Box plots depicting transcriptional correlation of clustered fd-BP pairs assigned to three groups based on the presence and clustering of TFs. ‘No TF’ refers to clustered fd- BP pairs in which one or both of the two BP terms is missing a TF (N=13) ; ‘No clustered TF’ refers to clustered fd-BP pairs in which both BP terms have a TF, but their TFs are not clustered (N=123); and ‘Clustered TF’ refers to clustered BP pairs in which both fd- BP pairs have a TF and share at least one clustered TF pair (N=142). *p*-values were calculated by the two-sample t-test. (C) Ring plot of human genome, drawn to the same specification as Figure 2E, depicting the transcriptional correlation of a representative clustered fd-BP pair with clustered TFs. Green and blue curves depict high transcriptional correlation (>0.5) between a clustered TF and genes in the other BP. The two BP terms are GO:0006959 humoral immune response (blue) and GO:0007162 negative regulation of cell adhesion (green) in human. (D) Graph depicting the transcriptional correlation of all human fd-BP pairs as a function of the strength of their genomic clustering. Red and blue lines represent transcriptional correlation of fd-BP pairs with (N=59,760) or without clustered TFs (N=401,118), respectively. BP pairs were binned by the strength of their gene clustering. The correlation plotted represents the moving average of 5,000 points.

The strong predictive value of clustered TFs in BP-BP correlation suggested that clustered TFs alone (without pathway-wide BP-BP clustering) could be sufficient to predict the transcriptional correlation of BP pairs. If true, this type of gene placement could couple the transcription of multiple fd-BP pairs. To test for this, we compared the transcriptional correlation of fd-BP pairs that share (N=58,760) or do not share (N=401,118) at least one clustered TF pair. By plotting BP-BP transcriptional correlation versus the strength of clustering, we found that (1) genome clustering positively correlates with BP-BP transcriptional correlation, and that (2) fd-BP pairs with at least one clustered TF have a dramatically higher transcriptional correlation compared to fd-BP pairs without a clustered TF (Figure 4D). In fact, we also found that the most concordant and discordant BP pairs have the highest TF-TF transcriptional correlation consistent with an activator-activator and activator-repressor TF pairing, respectively (Figure S10F). Likewise, fs-BP pairs (Lin Score >0.1) with at least one clustered TF (N=27,898) have a dramatically higher transcriptional correlation compared to fs-BP pairs without a clustered TF (N=158,016) (Figure S10E). In sum, these analyses demonstrate that beyond pathway-wide BP-BP gene clustering, TF clusters alone are strong predictors of the transcriptional coupling of their associated BPs.

### Analyses of biological pathway pairs defined by other platforms (KEGG, BioCarta, Wiki Pathways) confirm that clustered pathways, especially those with a clustered TF, are transcriptionally correlated in human cells

As mentioned above, we used the Gene Ontology database to define biological pathways in the five model organisms under study, providing a pan-organismal platform to assess the conservation of BP gene clustering across evolution. We further found that clustered BPs, especially those with a clustered TF pair, are transcriptionally correlated in human cells (Figure 4). Because hierarchical organization of Gene Ontology could introduce biases in pathway definition, here to assess whether biological pathways defined by other databases also produce similar results, we used three different commonly used human databases for defining biological pathways (KEGG, BioCarta, and WikiPathways)^91^ to assess the consistency of our findings across databases.

Because there is no equivalent measure (semantic similarity) to quantify similarities between pathway pairs on these platforms, we defined ‘disparate pathway pairs’ as those which do not share any overlapping genes. To assess whether this strategy enriches for disparate pathway pairs, we applied it to Gene Ontology BPs and found that BP pairs which share no overlapping genes have a lower average Lin Score (74% have a Lin <0.1) and display lower transcriptional correlation relative to those BP pairs which possess at least one overlapping gene (Figure S11). These together suggest that this approach enriches for disparate pathway pairs on these platforms.

Focusing on disparate pathway pairs, we found that, similar to our analyses with GO (Figure 1C-H), clustered pathway pairs can be identified on KEGG, BioCarta and WikiPathways platforms using the same statistical pipeline (Figure 1A-C) developed for GO analyses (Figure S12). Again, the QQ plots for pairwise clustering analyses on KEGG/BioCarta/Wiki-pathways platforms (Figure S12B-D) curl upwards indicating that the statistical platform can identify significantly clustered disparate pathway pairs in a database-independent manner (Figure 1D-H and Figure S4). In the end, the top 10% of the most significantly clustered (lowest *p*-value) disparate pathway pairs were selected (KEGG (N=1,349), BioCarta (N=3,454), and WikiPathways (N=14,773), and separated into three bins bases on the presence of clustered TFs (Figure S12C-D). Because clustered gene pairs tend to be transcriptionally correlated (Figure S1), these genes were excluded from all transcriptional correlation calculations. This permitted us to identify functionally disparate pathway pairs whose transcriptional correlations are pathway-wide, and not caused by a coincidental correlation of their clustered gene pairs. By calculating their transcriptional correlation across the 53 tissues on GTEx, we found that disparate clustered pathway pairs, especially those with a clustered TF pair, are transcriptionally correlated in human cells (Figure S13). These analyses are similar to those with GO BP terms (Figure 4), which together demonstrate that our findings are not restricted to a given database and its definition of biological pathways. They also demonstrate that beyond pathway-pathway clustering, TF clusters alone are strong predictors of the transcriptional coupling of their associated pathways, suggesting that TF clusters could function to link transcription between disparate pathway pairs.

### TF clustering is required for transcriptional coregulation of disparate BPs

If clustered TFs couple the transcription of their associated BPs, then their disruption (e.g. by a deletion or translocation) should result in the loss of these transcriptional couplings. To test this prediction, we used the Cancer Cell Line Encyclopedia (CCLE) dataset^92^, to identify cell lines in which a deletion or a translocation disrupts a clustered TF pair whose associated BPs are transcriptionally correlated. We then quantified the extent to which such a deletion or translocation impacts the transcriptional correlation of the associated BPs (see Methods). As an example, among the CCLE’s lymphocytic cell line collection, SUDHL8 is a cell line that carries a deletion in the IRF8 gene disrupting the IRF8-FOXF1 clustered TF pair. Comparing the SUDHL8 transcriptome versus the other lymphocytic cell lines by using standardized residuals as a measure for transcriptional deviation, we found that even though the overall transcriptome of SUDHL8 is typical for a subset of lymphocytic cell line (Figure 5A), specifically the transcriptional coherence among the IRF8- and FOXF1-associated BPs is lost (Figure 5B-C). Similarly, among the transverse colon cancer cell lines, the SNU1033 cell line carries a deletion in the PRDM16 gene, disrupting the PRDM16-TP73 clustered TF pair. Here also we observed that even though the overall transcriptome of SNU1033 is typical for a transverse colon cancer cell line (Figure S14A), specifically the transcriptional coherence among the PRDM16- andTP73- associated BPs is lost (Figure S14B-C).

**Figure 5.**
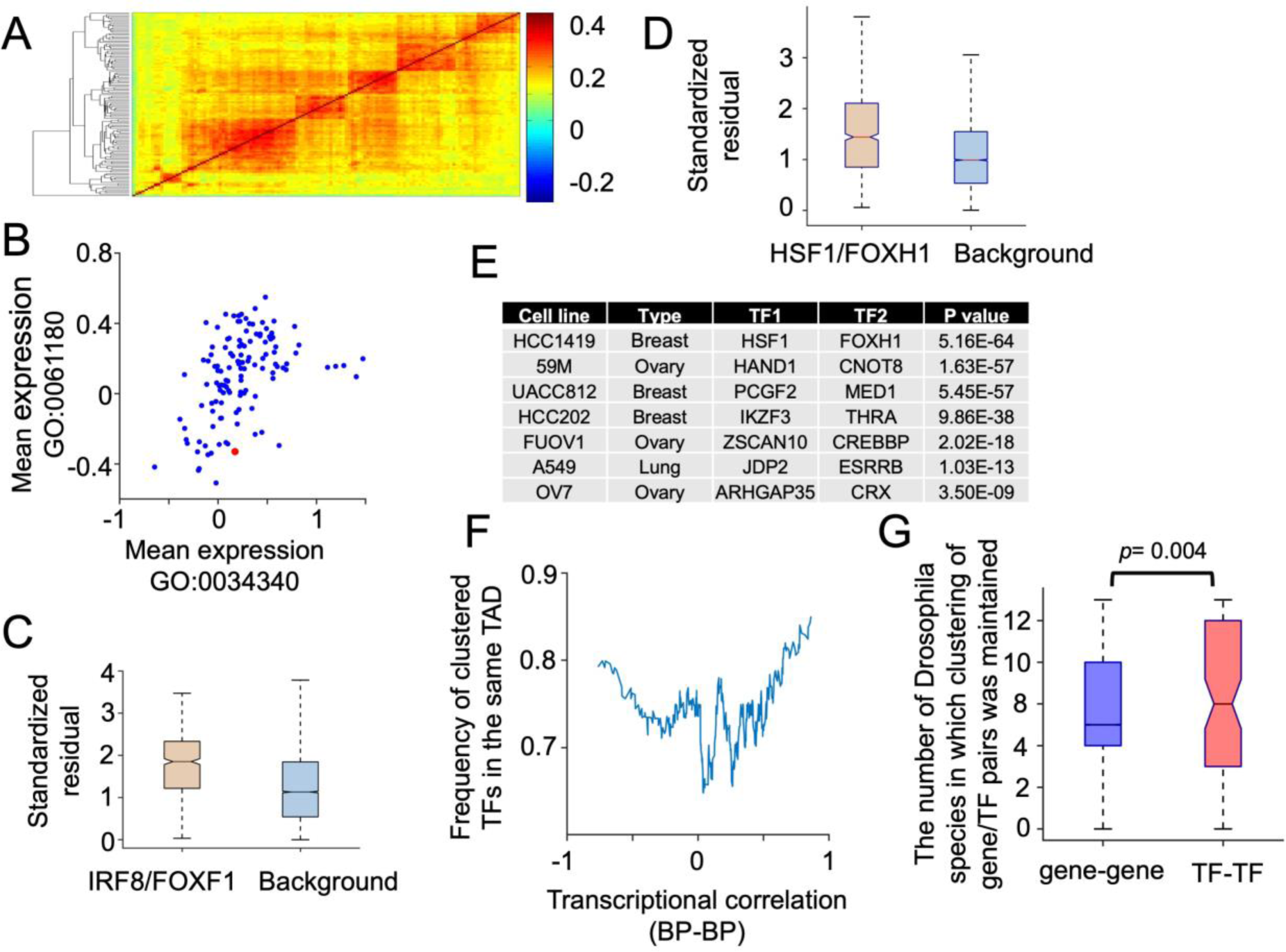
TF-TF clustering predicts the transcriptional coupling of the associated BP pairs and is maintained in evolution. (A) Hierarchical clustering of transcriptomes of EBV-transformed lymphocytic cancer cell lines (N=126). Red arrow indicates SUDHL8, the cell line with IRF8 deletion. The color indicates the overall pairwise correlation between two cancer cell lines. (B) Graph depicting the mean expression of two IRF8- and FOXF1- associated BPs (GO:0034340 response to type I interferon and GO:0061180 mammary gland epithelium development, respectively) among lymphocytic cell lines. Red dot represents SUDHL8. (C) Boxplot depicting the distribution of standardized residuals of BP pairs in SUDHL8 versus other lymphocytic cancer cell lines. The left and right bars depict this distribution for the IRF8/FOXF1-associated BP pairs and random (background) BP pairs, respectively. (D) Boxplot depicting the distribution of the standardized residuals of BP pairs in HCC1419 cell line, carrying a translocation disrupting HSF1-FOXH1 genomic clustering, versus other breast cancer cell lines. The left and right bars depict this distribution for the HSF1/FOXH1-associated BP pairs and 10,000 random BP pairs, respectively. (E) Examples of cancer cell lines where the occurrence of a translocation, physically separating a clustered TF pair, co-occurs with greater standard residuals of the BPs associated with the TFs versus random (background) BP pairs. Similar to C and D, standard residual of translocation-bearing cell lines was compared against other cancer cell lines from the same tissue. *p*-values are from two-sample *t*-tests. (F) Plot depicting the frequency that clustered TFs occur in the same TAD versus the transcriptional correlation of the associated fd-BPs. For each clustered TF pair, the mean correlation of associated fd-BP pairs was plotted. The frequency of TF occurring in the same TAD is the moving average of 50 points. (G) Graph comparing the maintenance of gene-gene versus TF-TF clustering among 12 Drosophila species. TF or gene pairs which are separated by less than 4X mean intergenic distance in the *D*. *melanogaster* genome were considered clustered. We then asked whether the distance between the corresponding TF or gene pair in the other Drosophila species was also less than 4X mean intergenic distance. *p*-values were calculated by the two-sample t-test.

Deletion of a TF could impact the expression of many genes in its pathway. To test if genic proximity of TFs specifically can predict the transcriptional coherence of their associated BPs, we analyzed the transcriptomes of cell lines carrying translocations between clustered TFs in a similar fashion to Figure 5A-C. Figure 5D illustrates one example where a translocation in the HCC1419 breast cancer cell line disrupts the clustering of HSF1-FOXH1 TF pair. Consistent with our model, we found that even though the overall transcriptome of this cell line is similar to other breast cancer lines, there is a specific loss of transcriptional coherence among the HSF1/FOXH1-associated BP pairs (Figure 5D). Figure 5E and Table S4 summarize similar findings in 31 other cancer cell lines from different tissues of origin in which a translocation separating a clustered TF pair co-occurs specifically with a significant loss of transcriptional correlation of their associated BPs. Taken together, these data support a model in which clustered TFs couple the transcriptional output of fd-BPs.

Because genes within TADs are transcriptionally correlated, another predication of our model is that the cooccurrence of the clustered TFs in the same TAD would portend the transcriptional correlations of their associated BPs. To test this prediction (see StarMethods), we analyzed TAD domains identified from 37 independent samples^59,93^ (Figure S15A) and plotted the frequency with which a clustered TF is found in the same TAD versus the transcriptional correlation of their associated BPs. As expected, clustered TFs found most frequently in the same TAD displayed the highest TF-TF transcriptional correlations (Figure S15B). Moreover, their associated BP pairs displayed the most highly concordant (activator-activator) and discordant (activator-repressor) transcriptional couplings (Figure 5F). Together these data further support a model in which transcriptionally coupled clustered TFs link the transcriptional outputs of their associated BPs in the human genome.

### Maintenance of TF-TF clusters among 12 *Drosophila* species

Our data suggest that clustered TFs play a critical role in coordinating the transcriptional outputs of fd-BPs. If so, then a prediction of this model is that maintenance of clustered TFs would be under selection considering TF-TF clusters could play an important role in establishing essential GRNs. To test this prediction, we examined the genomes of 12 Drosophila species (Figure S16) and asked whether TF-TF clustering is maintained preferentially relative to the clustering of non-TF gene pairs during the evolution of these Drosophila species. Consistent with our model, Figure 5G shows that clustered TF pairs are maintained more frequently compared to clustered non-TF gene pairs. Overall, these analyses together suggest that TF-TF clusters play important regulatory roles by linking the transcriptional output of fd-BPs, thus creating GRNs in eukaryotes.

## Discussion

Here, the analysis of the distantly related genome of yeast, fly, worm, mouse and human reveals that clustering of disparate BPs is a conserved feature of eukaryotic genomes, demonstrating a strong evolutionary selection for maintaining this type of genome organization. Clustered BP pairs display strong transcriptional correlation in human cells, especially those which also share a clustered TF. Moreover, TF clustering alone is a strong predictor of BP-BP transcriptional correlation, suggesting that formation and maintenance of TF clusters (which we propose form regulatory nodes) in evolution provide an efficient genomic architecture for coupling the transcription of GRN sub- circuits. In support of this, we find that deletion or translocation of a TF within a clustered TF pair impacts the transcriptional coherence of their associated BPs. Moreover, we show that TF clusters are maintained more frequently than gene-gene clusters, supporting their proposed critical role in acting as regulatory nodes for coupling the transcription of disparate BPs. Figure 6 depicts a simple model in which transcriptional coregulation of clustered TFs drive the transcriptional coherence of their respective BPs in *trans*. According to our model, eukaryotic genomes contain many regulatory nodes through which the combinatorial activities of *trans*-acting factors, such as TFs, can be efficiently coordinated. This in turn endows organisms with the ability to coregulate several GRN sub-circuits simultaneously, which could help the timely establishment of distinct biological outcomes rapidly and robustly (Figure 6).

**Figure 6.**
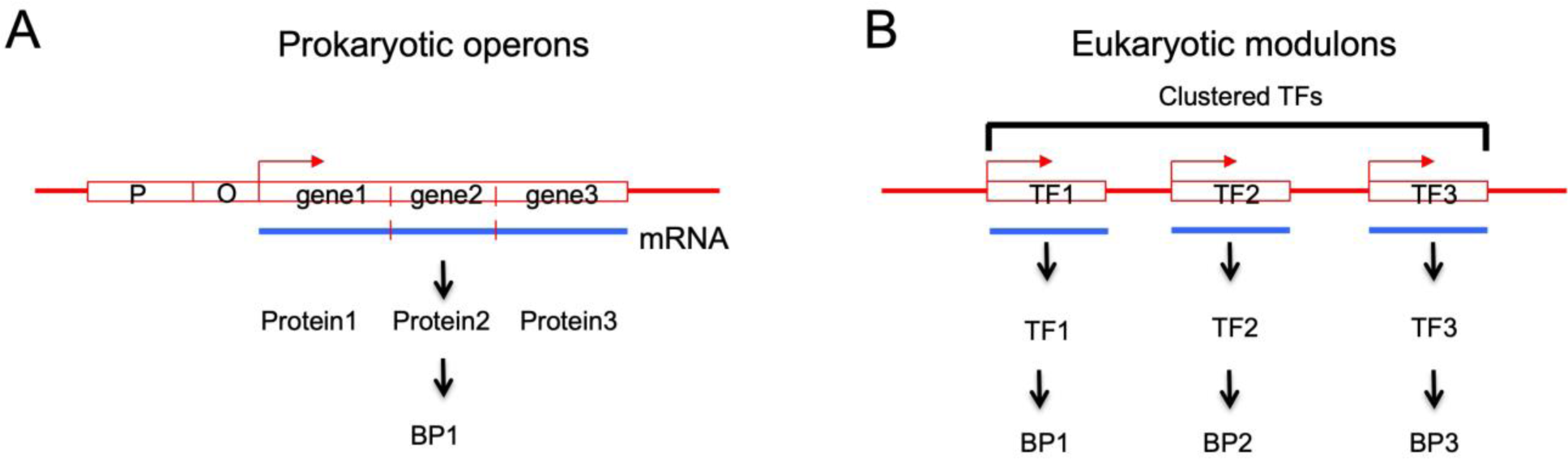
A model for how clustered TFs could help create modulons in eukaryotes. (A-B) Schematic model of (A) prokaryotic operons and (B) eukaryotic modulons regulated by clustered TFs. According to this model, transcriptional activation of clustered TFs can coordinate the expression of hundreds of genes belonging to fd-BPs dispersed throughout the genome.

### BP clustering across evolution

Previously, it was shown that genomic clustering of similarly expressed or functionally interrelated genes provides an efficient strategy for coregulating or temporally ordering the expression of many genes in the same pathway^27,35,45–48,69,72,94^. This helps in the establishment of transient, persistent or temporally ordered adaptive GRN states^95^, and is under strong evolutionary selection^11–13^. Accordingly, in support of this, a recent evolutionary survey of 341 fungal species revealed that evolution by vertical and horizontal gene transfer of several metabolic BPs has led to the maintenance of the same gene clusters among these fungal species. Strikingly, this was true even in cases of convergent evolution such that the *de novo* acquirement of some BPs leads to the formation of the same gene clusters^96^, demonstrating the importance of clustering of interrelated genes in evolution. Here in this report, we show that genomic clustering also extends to BP pairs (as defined by GO (Figure 1C-H) or other platforms (Figure S12)), many of which in normal human tissue display strong transcriptional correlation (Figure 4A, Figure S13A and Table S2). This correlation is pathway-wide and especially strong among BP pairs which also share a clustered TF pair (Figure 4B and Figure S13A). Moreover, even though individual genes, TFs and lower-level GRN sub-circuits vary across evolution, our data demonstrate that the maintenance of this type of genome organization for higher level BP-BP coregulation spans hundreds of millions of years of evolutionary time (Figure 3). In addition to revealing clustering among previously known interdependent BPs (e.g. stress response and metabolism BP pair, and amino acid metabolism and neural development BP pair), some clustered BP pairs illustrate emerging interdependencies. For example, we found that learning and histone methylation and learning and RNA methylation are clustered in worm and fly genomes, respectively (Table S3). Indeed, several recent reports have provided support for the emerging role of these modifications in learning and memory in metazoans^97–100^. Another example is the clustering of sphingolipid metabolism genes with serine/threonine kinase signaling genes in mouse and human genomes. Here also, recent data demonstrate the emerging role of sphingolipid metabolites in intra- and intercellular signaling pathways^101–103^, which when combined with their genome clustering with serine/threonine kinase signaling pathways suggest a functional link for their transcriptional coupling. Lastly, we also found that some BP clusters do not have clearly defined links in biology (Table S2). It will be informative to determine whether these kinds of clustered BP pairs portend previously unappreciated biological interdependencies considering their correlated transcriptional outputs in human cells.

Beyond pathway-wide BP-BP gene clustering, we also found that a strong predictor of transcriptional correlation between disparate BPs is the presence of at least one shared clustered TF pair (Figure 4D and Figure S10C), suggesting that clustered TFs can be used to couple the transcription of associated BPs. In support of this, we find that deletion or translocation of a TF within a clustered TF pair disrupt the transcriptional coherence of their associated BPs. (Figure 4 and Figure S14). Considering that during the continual shuffling of genes in evolution the probability of forming clustered TFs (a single gene pair) is higher than clustered BP pairs (several gene pairs), clustered TFs may present an efficient evolutionary solution for coregulation of disparate BPs and creation of tunable modulons. If so, then such a model would predict that the maintenance of TF clusters would be under strong evolutionary selection. In support of this, we found that the maintenance of TF-clusters is under stronger selective pressure as evidence by the preferential maintenance of TF-TF clusters over non-TF gene pairs (Figure 5G). This is consistent with the recent observations that combinations of TFs^104–106^ and enhancers^107^ define the various cell states in mammals. Because in the human genome, gene clusters showing the highest transcriptional coupling across different tissues occur in regions of high gene density^108^, we speculate that TF clusters in high gene density regions present promising targets for probing novel biological interrelationships between BPs.

In addition to protein-coding genes, clustering and coregulation of *trans*-acting noncoding factors, such as micro RNAs (miRNAs) also may play important roles in coregulation of multiple BPs, consistent with the presence of their target sequences in multiple transcripts^109^. These, together with TFs, provide additional layers of combinatorial regulation through which system-level changes to transcriptomes can be achieved. For example, TF-miRNA and miRNA-miRNA clustering also may serve as efficient strategies to regulate expression of a wide range of target genes. With better classification and identification of ncRNAs and their targets, the application of our clustering framework may expose novel BP interconnections and other networks motifs involving various coding and noncoding elements.

## Conclusions

Overall, based on these analyses we hypothesize that as eukaryotic genomes expanded in size and complexity during evolution, modulons were maintained and created by the formation of *trans*-acting regulatory nodes (such as TF clusters) through which the biological outputs of disparate BPs were coupled. This, we propose, is a conserved organizing principle of eukaryotic genomes. Additionally, the transcriptionally linked disparate BP pairs identified in this study not only support the well-known interdependencies among some disparate BPs, but also suggest the existence of new BP interconnections which future studies may uncover their molecular links. Finally, because the functional, hierarchical and temporal organization of these sub-circuits underlie the complex developmental and adaptive programs deployed in eukaryotes, in future studies it will be informative to ask whether and how the loss or inappropriate gain of these coregulations impacts disease states such as cancer.

## Acknowledgement

This work was supported by NIH (R01GM125782), American Cancer Society Research Scholar Award, the Harvard Ludwig Fund and Howard M. Goodman Fellowship to MM. We thank Meeta Mistry for her help with the semantic similarity analysis. We are grateful to Russell Jenkins for helpful suggestion regarding sphingolipid metabolism and kinase signaling. We also thank the members of the Motamedi Lab and Nick Dyson for critical reading of this manuscript.

## Author contribution

MM conceived the concepts and ideas. RJ developed the statistical framework in consultation with ML and MA. RJ performed all data analyses. ML and MA vetted the analyses. RJ and MM wrote the manuscript which was edited by ML and MA.

## Supplementary information

**Additional file 1: Supplemental Figures S1-16**.

**Additional file 2: Table S1**: List of all BP terms

**Additional file 3: Table S2**: List of clustered BP pairs, including the number of observed gene pairs and background, *p* and FDR values

**Additional file 4: Table S3**: Groups of BP terms based on hierarchical clustering

**Additional file 5: Table S4**: List of translocations between clustered TFs in CCLE collection which co-occur with the loss of transcriptional correlation of the associated BPs.

## Reference

1. Davidson, E.H., Rast, J.P., Oliveri, P., Ransick, A., Calestani, C., Yuh, C.H., Minokawa, T., Amore, G., Hinman, V., Arenas-Mena, C., et al. (2002). A genomic regulatory network for development. Science 295, 1669–1678. 10.1126/science.1069883.

2. Lee, T.I., Rinaldi, N.J., Robert, F., Odom, D.T., Bar-Joseph, Z., Gerber, G.K., Hannett, N.M., Harbison, C.T., Thompson, C.M., Simon, I., et al. (2002). Transcriptional regulatory networks in Saccharomyces cerevisiae. Science 298, 799–804. 10.1126/science.1075090.

3. Stathopoulos, A., and Levine, M. (2005). Genomic regulatory networks and animal development. Dev. Cell 9, 449–462. 10.1016/j.devcel.2005.09.005.

4. Tavazoie, S., Hughes, J.D., Campbell, M.J., Cho, R.J., and Church, G.M. (1999). Systematic determination of genetic network architecture. Nat. Genet. 22, 281–285. 10.1038/10343.

5. Friedman, N., Linial, M., Nachman, I., and Pe’er, D. (2000). Using Bayesian networks to analyze expression data. J. Comput. Biol. 7, 601–620. 10.1089/106652700750050961.

6. Vogelstein, B., Lane, D., and Levine, A.J. (2000). Surfing the p53 network. Nature 408, 307–310. 10.1038/35042675.

7. Erwin, D.H., and Davidson, E.H. (2009). The evolution of hierarchical gene regulatory networks. Nat. Rev. Genet. 10, 141–148. 10.1038/nrg2499.

8. Herrero, J., Valencia, A., and Dopazo, J. (2001). A hierarchical unsupervised growing neural network for clustering gene expression patterns. Bioinformatics 17, 126–136. 10.1093/bioinformatics/17.2.126.

9. Milo, R., Shen-Orr, S., Itzkovitz, S., Kashtan, N., Chklovskii, D., and Alon, U. (2002). Network motifs: simple building blocks of complex networks. Science 298, 824– 827.

10. Shen-Orr, S.S., Milo, R., Mangan, S., and Alon, U. (2002). Network motifs in the transcriptional regulation network of Escherichia coli. Nat. Genet. 31, 64–68. 10.1038/ng881.

11. Thompson, D., Regev, A., and Roy, S. (2015). Comparative analysis of gene regulatory networks: From network reconstruction to evolution. Annu. Rev. Cell Dev. Biol. 31, 399–428. 10.1146/annurev-cellbio-100913-012908.

12. Halfon, M.S. (2017). Perspectives on gene regulatory network evolution. Trends Genet. 33, 436–447. 10.1016/j.tig.2017.04.005.

13. Peter, I.S., and Davidson, E.H. (2011). Evolution of gene regulatory networks controlling body plan development. Cell 144, 970–985. 10.1016/j.cell.2011.02.017.

14. Maeso, I., Irimia, M., Tena, J.J., Casares, F., and Gómez-Skarmeta, J.L. (2013). Deep conservation of cis-regulatory elements in metazoans. Philos. Trans. R. Soc. B Biol. Sci. 368. 10.1098/rstb.2013.0020.

15. Jacob, F., and Monod, J. (1961). Genetic regulatory mechanisms in the synthesis of proteins. J. Mol. Biol. 3, 318–356. 10.1016/S0022-2836(61)80072-7.

16. Spieth, J., Brooke, G., Kuersten, S., Lea, K., and Blumenthal, T. (1993). Operons in C. elegans: polycistronic mRNA precursors are processed by trans-splicing of SL2 to downstream coding regions. Cell 73, 521–532.

17. Brogna, S., and Ashburner, M. (1997). The Adh-related gene of Drosophila melanogaster is expressed as a functional dicistronic messenger RNA: multigenic transcription in higher organisms. EMBO J. 16, 2023–2031. 10.1093/emboj/16.8.2023.

18. Blumenthal, T., Davis, P., and Garrido-Lecca, A. (2015). Operon and non-operon gene clusters in the C. elegans genome. WormBook Online Rev. C Elegans Biol., 1–20. 10.1895/wormbook.1.175.1.

19. Crosby, M.A., Sian Gramates, L., dos Santos, G., Matthews, B.B., St. Pierre, S.E., Zhou, P., Schroeder, A.J., Falls, K., Emmert, D.B., Russo, S.M., et al. (2015). Gene model annotations for Drosophila melanogaster: The rule-benders. G3 Genes Genomes Genet. 5, 1737–1749. 10.1534/g3.115.018937.

20. Michalak, K., Orr, W.C., and Radyuk, S.N. (2008). Drosophila peroxiredoxin 5 is the second gene in a dicistronic operon. Biochem. Biophys. Res. Commun. 368, 273–278. 10.1016/j.bbrc.2008.01.052.

21. Misra, S., Crosby, M.A., Mungall, C.J., Matthews, B.B., Campbell, K.S., Hradecky, P., Huang, Y., Kaminker, J.S., Millburn, G.H., Prochnik, S.E., et al. (2002). Annotation of the Drosophila melanogaster euchromatic genome: a systematic review. Genome Biol. 3, RESEARCH0083. 10.1186/gb-2002-3-12-research0083.

22. Cohen, B.A., Pilpel, Y., Mitra, R.D., and Church, G.M. (2002). Discrimination between paralogs using microarray analysis: application to the Yap1p and Yap2p transcriptional networks. Mol. Biol. Cell 13, 1608–1614. 10.1091/mbc.01-10-0472.

23. Papp, B., Pál, C., and Hurst, L.D. (2003). Dosage sensitivity and the evolution of gene families in yeast. Nature 424, 194–197. 10.1038/nature01771.

24. Poyatos, J.F., and Hurst, L.D. (2007). The determinants of gene order conservation in yeasts. Genome Biol. 8, R233. 10.1186/gb-2007-8-11-r233.

25. Spellman, P.T., and Rubin, G.M. (2002). Evidence for large domains of similarly expressed genes in the Drosophila genome. J. Biol. 1, 5.

26. Mezey, J.G., Nuzhdin, S. V, Ye, F., and Jones, C.D. (2008). Coordinated evolution of co-expressed gene clusters in the Drosophila transcriptome. BMC Evol. Biol. 8, 2. 10.1186/1471-2148-8-2.

27. Weber, C.C., and Hurst, L.D. (2011). Support for multiple classes of local expression clusters in Drosophila melanogaster, but no evidence for gene order conservation. Genome Biol. 12, R23. 10.1186/gb-2011-12-3-r23.

28. Blumenthal, T., and Gleason, K.S. (2003). Caenorhabditis elegans operons: form and function. Nat. Rev. Genet. 4, 110–118. 10.1038/nrg995.

29. Roy, P.J., Stuart, J.M., Lund, J., and Kim, S.K. (2002). Chromosomal clustering of muscle-expressed genes in Caenorhabditis elegans. Nature 418, 975–979. 10.1038/nature01012.

30. Kamath, R.S., Fraser, A.G., Dong, Y., Poulin, G., Durbin, R., Gotta, M., Kanapin, A., Le Bot, N., Moreno, S., Sohrmann, M., et al. (2003). Systematic functional analysis of the Caenorhabditis elegans genome using RNAi. Nature 421, 231–237. 10.1038/nature01278.

31. Tsai, H.K., Huang, P.Y., Kao, C.Y., and Wang, D. (2009). Co-expression of neighboring genes in the zebrafish (Danio rerio) genome. Int. J. Mol. Sci. 10, 3658– 3670. 10.3390/ijms10083658.

32. Ng, Y.K., Wu, W., and Zhang, L. (2009). Positive correlation between gene coexpression and positional clustering in the zebrafish genome. BMC Genomics 10, 42. 10.1186/1471-2164-10-42.

33. Li, Q., Lee, B.T., and Zhang, L. (2005). Genome-scale analysis of positional clustering of mouse testis-specific genes. BMC Genomics 6, 7. 10.1186/1471-2164-6-7.

34. Singer, G.A.C., Lloyd, A.T., Huminiecki, L.B., and Wolfe, K.H. (2005). Clusters of co-expressed genes in mammalian genomes are conserved by natural selection. Mol. Biol. Evol. 22, 767–775. 10.1093/molbev/msi062.

35. Caron, H., van Schaik, B., van der Mee, M., Baas, F., Riggins, G., van Sluis, P., Hermus, M.C., van Asperen, R., Boon, K., Voûte, P.A., et al. (2001). The human transcriptome map: clustering of highly expressed genes in chromosomal domains. Science 291, 1289–1292. 10.1126/science.1056794.

36. Lee, J.M., and Sonnhammer, E.L.L. (2003). Genomic gene clustering analysis of pathways in eukaryotes. Genome Res. 13, 875–882. 10.1101/gr.737703.

37. Fukuoka, Y., Inaoka, H., and Kohane, I.S. (2004). Inter-species differences of co-expression of neighboring genes in eukaryotic genomes. BMC Genomics 5, 4. 10.1186/1471-2164-5-4.

38. Sémon, M., Lobry, J.R., and Duret, L. (2006). No evidence for tissue-specific adaptation of synonymous codon usage in humans. Mol. Biol. Evol. 23, 523–529. 10.1093/molbev/msj053.

39. Makino, T., and McLysaght, A. (2008). Interacting gene clusters and the evolution of the vertebrate immune system. Mol. Biol. Evol. 25, 1855–1862. 10.1093/molbev/msn137.

40. Michalak, P. (2008). Coexpression, coregulation, and cofunctionality of neighboring genes in eukaryotic genomes. Genomics 91, 243–248. 10.1016/J.YGENO.2007.11.002.

41. Al-Shahrour, F., Minguez, P., Marqués-Bonet, T., Gazave, E., Navarro, A., and Dopazo, J. (2010). Selection upon genome architecture: conservation of functional neighborhoods with changing genes. PLoS Comput. Biol. 6, e1000953. 10.1371/journal.pcbi.1000953.

42. Ebisuya, M., Yamamoto, T., Nakajima, M., and Nishida, E. (2008). Ripples from neighbouring transcription. Nat. Cell Biol. 10, 1106–1113. 10.1038/ncb1771.

43. Vandepoele, K., Quimbaya, M., Casneuf, T., De Veylder, L., and Van de Peer, Y. (2009). Unraveling transcriptional control in Arabidopsis using cis-regulatory elements and coexpression networks. Plant Physiol. 150, 535–546. 10.1104/pp.109.136028.

44. Feuerborn, A., and Cook, P.R. (2015). Why the activity of a gene depends on its neighbors. Trends Genet. TIG 31, 483–490. 10.1016/j.tig.2015.07.001.

45. Lercher, M.J., Urrutia, A.O., and Hurst, L.D. (2002). Clustering of housekeeping genes provides a unified model of gene order in the human genome. Nat. Genet. 31, 180–183. 10.1038/ng887.

46. Amores, A., Force, A., Yan, Y.L., Joly, L., Amemiya, C., Fritz, A., Ho, R.K., Langeland, J., Prince, V., Wang, Y.L., et al. (1998). Zebrafish hox clusters and vertebrate genome evolution. Science 282, 1711–1714.

47. Lemons, D., and McGinnis, W. (2006). Genomic evolution of Hox gene clusters. Science 313, 1918–1922. 10.1126/science.1132040.

48. Versteeg, R., van Schaik, B.D.C., van Batenburg, M.F., Roos, M., Monajemi, R., Caron, H., Bussemaker, H.J., and van Kampen, A.H.C. (2003). The human transcriptome map reveals extremes in gene density, intron length, GC content, and repeat pattern for domains of highly and weakly expressed genes. Genome Res. 13, 1998–2004. 10.1101/gr.1649303.

49. Boulding, H., and Webber, C. (2012). Large-scale objective association of mouse phenotypes with human symptoms through structural variation identified in patients with developmental disorders. Hum. Mutat. 33, 874–883. 10.1002/humu.22069.

50. Doelken, S.C., Kohler, S., Mungall, C.J., Gkoutos, G. V., Ruef, B.J., Smith, C., Smedley, D., Bauer, S., Klopocki, E., Schofield, P.N., et al. (2013). Phenotypic overlap in the contribution of individual genes to CNV pathogenicity revealed by cross-species computational analysis of single-gene mutations in humans, mice and zebrafish. Dis. Model. Mech. 6, 358–372. 10.1242/dmm.010322.

51. Golzio, C., Willer, J., Talkowski, M.E., Oh, E.C., Taniguchi, Y., Jacquemont, S., Reymond, A., Sun, M., Sawa, A., Gusella, J.F., et al. (2012). KCTD13 is a major driver of mirrored neuroanatomical phenotypes of the 16p11.2 copy number variant. Nature 485, 363–367. 10.1038/nature11091.

52. Andrews, T., Honti, F., Pfundt, R., de Leeuw, N., Hehir-Kwa, J., Vulto-van Silfhout, A., de Vries, B., and Webber, C. (2015). The clustering of functionally related genes contributes to CNV-mediated disease. Genome Res. 25, 802–813. 10.1101/gr.184325.114.

53. Ben-Elazar, S., Yakhini, Z., and Yanai, I. (2013). Spatial localization of co-regulated genes exceeds genomic gene clustering in the Saccharomyces cerevisiae genome. Nucleic Acids Res. 41, 2191–2201. 10.1093/nar/gks1360.

54. Ben-Elazar, S., Chor, B., and Yakhini, Z. (2019). The Functional 3D Organization of Unicellular Genomes. Sci. Rep. 9, 12734. 10.1038/s41598-019-48798-7.

55. Diament, A., and Tuller, T. (2017). Tracking the evolution of 3D gene organization demonstrates its connection to phenotypic divergence. Nucleic Acids Res. 45, 4330–4343. 10.1093/nar/gkx205.

56. Diament, A., Pinter, R.Y., and Tuller, T. (2014). Three-dimensional eukaryotic genomic organization is strongly correlated with codon usage expression and function. Nat. Commun. 5, 5876. 10.1038/ncomms6876.

57. Homouz, D., and Kudlicki, A.S. (2013). The 3D organization of the yeast genome correlates with co-expression and reflects functional relations between genes. PloS One 8, e54699. 10.1371/journal.pone.0054699.

58. Li, S., and Heermann, D.W. (2013). Transcriptional regulatory network shapes the genome structure of Saccharomyces cerevisiae. Nucl. Austin Tex 4, 216–228. 10.4161/nucl.24875.

59. Dixon, J.R., Selvaraj, S., Yue, F., Kim, A., Li, Y., Shen, Y., Hu, M., Liu, J.S., and Ren, B. (2012). Topological domains in mammalian genomes identified by analysis of chromatin interactions. Nature 485, 376–380. 10.1038/nature11082.

60. Lupiáñez, D.G., Kraft, K., Heinrich, V., Krawitz, P., Brancati, F., Klopocki, E., Horn, D., Kayserili, H., Opitz, J.M., Laxova, R., et al. (2015). Disruptions of topological chromatin domains cause pathogenic rewiring of gene-enhancer interactions. Cell 161, 1012–1025. 10.1016/j.cell.2015.04.004.

61. Lieberman-Aiden, E., van Berkum, N.L., Williams, L., Imakaev, M., Ragoczy, T., Telling, A., Amit, I., Lajoie, B.R., Sabo, P.J., Dorschner, M.O., et al. (2009). Comprehensive mapping of long-range interactions reveals folding principles of the human genome. Science 326, 289–293. 10.1126/science.1181369.

62. Mifsud, B., Tavares-Cadete, F., Young, A.N., Sugar, R., Schoenfelder, S., Ferreira, L., Wingett, S.W., Andrews, S., Grey, W., Ewels, P.A., et al. (2015). Mapping long-range promoter contacts in human cells with high-resolution capture Hi-C. Nat. Genet. 47, 598–606. 10.1038/ng.3286.

63. Le Dily, F., Baù, D., Pohl, A., Vicent, G.P., Serra, F., Soronellas, D., Castellano, G., Wright, R.H.G., Ballare, C., Filion, G., et al. (2014). Distinct structural transitions of chromatin topological domains correlate with coordinated hormone-induced gene regulation. Genes Dev. 28, 2151–2162. 10.1101/gad.241422.114.

64. Irimia, M., Maeso, I., Roy, S.W., and Fraser, H.B. (2013). Ancient cis-regulatory constraints and the evolution of genome architecture. Trends Genet. 29, 521–528. 10.1016/j.tig.2013.05.008.

65. Irimia, M., Tena, J.J., Alexis, M.S., Fernandez-Miñan, A., Maeso, I., Bogdanović, O., De La Calle-Mustienes, E., Roy, S.W., Gómez-Skarmeta, J.L., and Fraser, H.B. (2012). Extensive conservation of ancient microsynteny across metazoans due to cis-regulatory constraints. Genome Res. 22, 2356–2367. 10.1101/gr.139725.112.

66. Tan, L., Xing, D., Daley, N., and Xie, X.S. (2019). Three-dimensional genome structures of single sensory neurons in mouse visual and olfactory systems. Nat. Struct. Mol. Biol. 26, 297–307. 10.1038/s41594-019-0205-2.

67. Joh, R.I., Khanduja, J.S., Calvo, I.A., Mistry, M., Palmieri, C.M.C.M., Savol, A.J.A.J., Ho Sui, S.J.S.J., Sadreyev, R.I.R.I., Aryee, M.J.M.J., and Motamedi, M. (2016). Survival in quiescence requires the euchromatic deployment of Clr4/SUV39H by Argonaute-associated small RNAs. Mol. Cell 64, 1088–1101. 10.1016/j.molcel.2016.11.020.

68. Valcourt, J.R., Lemons, J.M.S., Haley, E.M., Kojima, M., Demuren, O.O., and Coller, H.A. (2012). Staying alive. 10.4161/cc.19879.

69. Cohen, B.A., Mitra, R.D., Hughes, J.D., and Church, G.M. (2000). A computational analysis of whole-genome expression data reveals chromosomal domains of gene expression. Nat. Genet. 26, 183–186. 10.1038/79896.

70. Janicki, S.M., Tsukamoto, T., Salghetti, S.E., Tansey, W.P., Sachidanandam, R., Prasanth, K. V, Ried, T., Shav-Tal, Y., Bertrand, E., Singer, R.H., et al. (2004). From silencing to gene expression: real-time analysis in single cells. Cell 116, 683–698. 10.1016/s0092-8674(04)00171-0.

71. Purmann, A., Toedling, J., Schueler, M., Carninci, P., Lehrach, H., Hayashizaki, Y., Huber, W., and Sperling, S. (2007). Genomic organization of transcriptomes in mammals: Coregulation and cofunctionality. Genomics 89, 580–587. 10.1016/j.ygeno.2007.01.010.

72. Eldabagh, R.S., Mejia, N.G., Barrett, R.L., Monzo, C.R., So, M.K., Foley, J.J., and Arnone, J.T. (2018). Systematic identification, characterization, and conservation of adjacent-gene coregulation in the budding yeast Saccharomyces cerevisiae. mSphere 3. 10.1128/mSphere.00220-18.

73. Hug, L.A., Baker, B.J., Anantharaman, K., Brown, C.T., Probst, A.J., Castelle, C.J., Butterfield, C.N., Hernsdorf, A.W., Amano, Y., Ise, K., et al. (2016). A new view of the tree of life. Nat. Microbiol. 1, 16048. 10.1038/nmicrobiol.2016.48.

74. Ciccarelli, F.D., Doerks, T., von Mering, C., Creevey, C.J., Snel, B., and Bork, P. (2006). Toward automatic reconstruction of a highly resolved tree of life. Science 311, 1283–1287. 10.1126/science.1123061.

75. Ashburner, M., Ball, C.A., Blake, J.A., Botstein, D., Butler, H., Cherry, J.M., Davis, A.P., Dolinski, K., Dwight, S.S., Eppig, J.T., et al. (2000). Gene ontology: tool for the unification of biology. Nat. Genet. 25, 25–29. 10.1038/75556.

76. Storey, J.D. (2003). The positive false discovery rate: a Bayesian interpretation and the q -value. Ann. Stat. 31, 2013–2035. 10.1214/aos/1074290335.

77. Mouse Genome Sequencing Consortium, Waterston, R.H., Lindblad-Toh, K., Birney, E., Rogers, J., Abril, J.F., Agarwal, P., Agarwala, R., Ainscough, R., Alexandersson, M., et al. (2002). Initial sequencing and comparative analysis of the mouse genome. Nature 420, 520–562. 10.1038/nature01262.

78. Pesquita, C. (2017). Semantic Similarity in the Gene Ontology. In The Gene Ontology Handbook Methods in Molecular Biology., C. Dessimoz and N. Škunca, eds. (Springer), pp. 161–173. 10.1007/978-1-4939-3743-1_12.

79. Lin, D. (1998). An information-theoretic definition of similarity. Proc. Fifteenth Int. Conf. Mach. Learn., 296–304.

80. Resnik, P. (1995). Using information content to evaluate semantic similarity in a taxonomy. Proc. 14TH Int. Jt. Conf. Artif. Intell. 1, 448--453.

81. Hotamisligil, G.S., and Davis, R.J. (2016). Cell signaling and stress responses. Cold Spring Harb. Perspect. Biol. 8, a006072. 10.1101/cshperspect.a006072.

82. Wellen, K.E., and Thompson, C.B. (2010). Cellular metabolic stress: Considering how cells respond to nutrient excess. Mol. Cell 40, 323–332. 10.1016/j.molcel.2010.10.004.

83. Green, D.R., Galluzzi, L., and Kroemer, G. (2014). Metabolic control of cell death. Science 345, 1250256–1250256. 10.1126/science.1250256.

84. Fernstrom, J.D., and Fernstrom, M.H. (2007). Tyrosine, phenylalanine, and catecholamine synthesis and function in the brain. J. Nutr. 137, 1539S–1547S. 10.1093/jn/137.6.1539s.

85. Fernstrom, J.D. (1990). Aromatic amino acids and monoamine synthesis in the central nervous system: influence of the diet. J. Nutr. Biochem. 1, 508–517. 10.1016/0955-2863(90)90033-H.

86. Jakeman, P.M. (1998). Amino acid metabolism, branched-chain amino acid feeding and brain monoamine function. Proc. Nutr. Soc. 57, 35–41. 10.1079/pns19980007.

87. Usuda, K., Kawase, T., Shigeno, Y., Fukuzawa, S., Fujii, K., Zhang, H., Tsukahara, T., Tomonaga, S., Watanabe, G., Jin, W., et al. (2018). Hippocampal metabolism of amino acids by L-amino acid oxidase is involved in fear learning and memory. Sci. Rep. 8, 11073. 10.1038/s41598-018-28885-x.

88. The GTEx Consortium (2013). The Genotype-Tissue Expression (GTEx) project. Nat. Genet. 45, 580–585. 10.1038/ng.2653.

89. Mao, L., Van Hemert, J.L., Dash, S., and Dickerson, J.A. (2009). Arabidopsis gene co-expression network and its functional modules. BMC Bioinformatics 10, 346. 10.1186/1471-2105-10-346.

90. Tornow, S., and Mewes, H.W. (2003). Functional modules by relating protein interaction networks and gene expression. Nucleic Acids Res. 31, 6283–6289. 10.1093/NAR/GKG838.

91. Liberzon, A., Subramanian, A., Pinchback, R., Thorvaldsdóttir, H., Tamayo, P., and Mesirov, J.P. (2011). Molecular signatures database (MSigDB) 3.0. Bioinformatics 27, 1739–1740. 10.1093/bioinformatics/btr260.

92. Barretina, J., Caponigro, G., Stransky, N., Venkatesan, K., Margolin, A.A., Kim, S., Wilson, C.J., Lehár, J., Kryukov, G. V., Sonkin, D., et al. (2012). The Cancer Cell Line Encyclopedia enables predictive modelling of anticancer drug sensitivity. Nature 483, 603–607. 10.1038/nature11003.

93. Wang, Y., Song, F., Zhang, B., Zhang, L., Xu, J., Kuang, D., Li, D., Choudhary, M.N.K., Li, Y., Hu, M., et al. (2018). The 3D genome browser: a web-based browser for visualizing 3D genome organization and long-range chromatin interactions. Genome Biol. 19, 151. 10.1186/s13059-018-1519-9.

94. Wang, G.-Z., Chen, W.-H., and Lercher, M.J. (2011). Coexpression of linked gene pairs persists long after their separation. Genome Biol. Evol. 3, 565–570. 10.1093/gbe/evr049.

95. Peter, I.S., and Davidson, E.H. (2016). Implications of developmental gene regulatory networks inside and outside developmental biology. Curr. Top. Dev. Biol. 117, 237–251. 10.1016/bs.ctdb.2015.12.014.

96. Marcet-Houben, M., and Gabaldón, T. (2019). Evolutionary and functional patterns of shared gene neighbourhood in fungi. Nat. Microbiol. 4, 2383–2392. 10.1038/s41564-019-0552-0.

97. Li, J., Yang, X., Qi, Z., Sang, Y., Liu, Y., Xu, B., Liu, W., Xu, Z., and Deng, Y. (2019). The role of mRNA m6A methylation in the nervous system. Cell Biosci. 9, 66. 10.1186/s13578-019-0330-y.

98. Widagdo, I.S., Pratt, N.L., and Roughead, E.E. (2018). The association between frailty and medicines use over time: an analysis using the Australian Longitudinal Study on Ageing population. J. Pharm. Pract. Res. 48, 405–415. 10.1002/jppr.1407.

99. Jarome, T.J., and Lubin, F.D. (2013). Histone lysine methylation: critical regulator of memory and behavior. Rev. Neurosci. 24, 375–387. 10.1515/revneuro-2013-0008.

100. Collins, B.E., Greer, C.B., Coleman, B.C., and Sweatt, J.D. (2019). Histone H3 lysine K4 methylation and its role in learning and memory. Epigenetics Chromatin 12, 7. 10.1186/s13072-018-0251-8.

101. Cartier, A., and Hla, T. (2019). Sphingosine 1-phosphate: Lipid signaling in pathology and therapy. Science 366, eaar5551. 10.1126/science.aar5551.

102. Montefusco, D.J., Matmati, N., and Hannun, Y.A. (2014). The yeast sphingolipid signaling landscape. Chem. Phys. Lipids 177, 26–40. 10.1016/j.chemphyslip.2013.10.006.

103. Hannun, Y.A., and Obeid, L.M. (2018). Sphingolipids and their metabolism in physiology and disease. Nat. Rev. Mol. Cell Biol. 19, 175–191. 10.1038/nrm.2017.107.

104. Braun, T., and Gautel, M. (2011). Transcriptional mechanisms regulating skeletal muscle differentiation, growth and homeostasis. Nat. Rev. Mol. Cell Biol. 12, 349–361. 10.1038/nrm3118.

105. Martello, G., and Smith, A. (2014). The nature of embryonic stem cells. Annu. Rev. Cell Dev. Biol. 30, 647–675. 10.1146/annurev-cellbio-100913-013116.

106. Waardenberg, A.J., Ramialison, M., Bouveret, R., and Harvey, R.P. (2014). Genetic networks governing heart development. Cold Spring Harb. Perspect. Med. 4, a013839. 10.1101/cshperspect.a013839.

107. Kieffer-Kwon KR, Tang Z, Mathe E, Qian J, Sung MH, Li G, Resch W, Baek S, Pruett N, Grøntved L, et al. (2013). Interactome maps of mouse gene regulatory domains reveal basic principles of transcriptional regulation. Cell 155, 1507–1520.

108. Ghanbarian, A.T., and Hurst, L.D. (2015). Neighboring genes show correlated evolution in gene expression. Mol. Biol. Evol. 32, 1748–1766. 10.1093/molbev/msv053.

109. Hausser, J., and Zavolan, M. (2014). Identification and consequences of miRNA– target interactions — beyond repression of gene expression. Nat. Rev. Genet. 15, 599– 612. 10.1038/nrg3765.

