## Supplementary material for "Gene clustering drives the transcriptional coherence of disparate biological processes in eukaryotes": Methods

### Supplementary Tables

**Table S1**: List of all BP terms

**Table S2**: List of clustered BP pairs, including the number of observed gene pairs and background, p and pFDR values

**Table S3**: Groups of BP terms based on hierarchical clustering

**Table S4**: List of translocations between clustered TFs in CCLE collection, which cooccur with the loss of transcriptional correlation of the associated BPs.

Key Resources Table

| REAGENT or RESOURCE | SOURCE | IDENTIFIER |
|---|---|---|
| Data Source | | |
| GO annotation and gene list for *S. pombe* | Pombase[1] | https://pombase.org |
| GO annotation file for *C. elegans*, *D. melanogaster*, *M. musculus* and *H. sapiens* | Gene Ontology Consortium[2,3] | https://www.geneontology.org/ |
| Gene list for *C. elegans*, *D. melanogaster*, *M. musculus* and *H. sapiens* | Ensemble Genomes[4] | https://ensemblgenomes.org/ |
| Human transcriptome | GTEx portal[5] | https://gtexportal.org/home/ |
| Transcriptome and copy number variation in cancer cell lines | Cancer Cell Line Encyclopedia (CCLE)[6] | https://portals.broadinstitute.org/ccle |
| List of human transcription factors | UniProt[7] | https://www.uniprot.org |
| Alternative genesets in human (KEGG, BioCarta, and Wiki pathways) | MSigDB[8,9] | https://www.gsea-msigdb.org/gsea/msigdb/ |
| Fly genomes | FlyBase[10] | https://flybase.org/ |
| Software and Algorithms | | |
| MATLAB R2017a/R2013a | MathWorks | https://www.mathworks.com/products/matlab.html |
| R v3.4.2 | R Core Team | http://www.R-project.org/ |
| GOSemSim v.0.5.5 | Bioconductor[11] | https://bioconductor.org/packages/release/bioc/html/GOSemSim.html |
| Circlize v.2.4.1 | Bioconductor[12] | https://cran.r-project.org/web/packages/circlize/index.html |
| TimeTree | TimeTree[13] | https://www.timetree.org/ |
| Deposited Data | | |

| | | |
|---|---|---|
| Code for analyses | This paper | https://figshare.com/projects/Transcriptional_coherence/72644 |

### Materials and Methods

#### Gene lists and gene ontology (GO) terms

The gene ontology (GO) annotation files for *S. pombe* and other organisms (*D. melanogaster*, *C. elegans*, *M. musculus*, and *H. sapiens*) were obtained from PomBase[1] and Gene Ontology Consortium[2,3], respectively. To focus our analyses on biological pathways, we chose Biological Process (BP) GO terms only. We also limited our analyses to BP GO terms with gene numbers suitable for generating sufficient statistical power in our analyses. In each organism, these gene number cutoffs were set after accounting for the total number of genes with at least one associated BP GO term and the number of chromosomes. For *S. pombe*, BP GO terms which contain10-250 genes, for *C. elegans* and *D. melanogaster* BPs with 20-250 genes, and for *M. musculus* and *H. sapiens* BPs with 50-250 genes were considered for analysis (for total BPs in each organism, see Figure 1C). The upper limit was used because BPs with many genes are the parent terms of several daughter BPs (with fewer genes), thus fail to capture specific biological functions. Here, we are interested in determining whether genomic clustering of specific, disparate BP pairs portend their transcriptional coregulation, thus we eliminated these broad terms from our analyses. Also, calculating background clustering (see below) produces statistically insignificant results. The list of genes and their coordinates are from PomBase[1] for *S. pombe* and Ensemble Genomes[4] for the other model organisms. We also removed largely redundant BP terms as defined by those containing 75% or more identical genes. In such instances, the BP term with more genes was retained and the other was eliminated from analyses. This reduced the number of BP-BP analyses between largely redundant BP pairs. The number of genes and BP terms analyzed in this work can be found in Figure 1C. The list of all BP terms in each organism can be found at https://figshare.com/projects/Transcriptional_coherence/72644.

#### Identification of clustered BP pairs

#### Rationale for selecting a threshold distance for gene-gene clustering

The primary reason for using a threshold distance for gene clustering in these analyses stemmed from the work in the fission yeast. Briefly, we found that when fission yeast cells enter quiescence (G0), many of the euchromatic genes which become co-regulated by heterochromatin factors were found in linear gene arrays with an average size of roughly 16-18Kb[14]. This led us to hypothesize that pathways whose genes need to be co-regulated transcriptionally under certain environmental or developmental conditions are maintained close to one another to create novel GRNs efficiently. In other words, instead of targeting hundreds of individual genes separately, targeting a few gene clusters containing these genes could provide an efficient means for their co-regulation. This hypothesis favored the use of a threshold distance for gene clustering that captured majority of the proximity-driven gene pair transcriptional correlations. Even though other methods genome clustering pathways have been developed which operate independently of a threshold distance[15], they are not ideally suited considering the biological basis of our hypothesis.

Because in most eukaryotes *cis*-regulatory elements and local gene-gene interactions driving transcriptional correlation of gene pairs tend to occur within five times (5X) the average intergenic distance[16,17], we assessed whether this distance (1MB in human) also captures the majority of proximity-driven gene pair transcriptional correlations found in the human genome. We used the human GTEx datasets[5] and quantified transcriptional correlation among all protein-coding gene pairs across different tissues (see below for details on how transcriptional correlations were calculated). Consistent with previous work[18,19], our data revealed that (1) transcriptional correlation is a function of the gene pair's intergenic distance, and (2) 5X mean intergenic distance (1Mb) is a good threshold distance capturing the majority of *cis* transcriptional correlations (Figure S1). Additionally, this distance corresponds to the average size of topologically associated domains (TADs) (or co-regulated domains) in this genome[20]. The threshold distance used in this study are 11,954bp for *S. pombe*, 45,613 for *C. elegans*, 61,141bp for *D. melanogaster*, 597,010bp for *M. musculus*, and 945,370bp for *H. sapiens*.

Consistent with the notion that local interactions driving transcriptional correlation of gene pairs tend to occur within five times (5X) the average intergenic distance, these threshold distances are a reasonable estimate for the average size of coregulated domains in other genomes. For example, the threshold distance for the fission yeast (12 kb), is roughly the same as the average linear gene array size (18 kb) we discovered in quiescent fission yeast cells[14]*.* Similarly, Drosophila melanogaster (60 kb) and mouse (600 kb) threshold distances are close to the average TAD size of 23-63kb and 880kb found in the Drosophila[21], and mouse[17] genomes. The only genome whose 5X mean intergenic distance lies outside of known co-regulated domains (e.g. TAD) is that of *C. elegans*; however, Figure S3 reveals that this distance captures a cross-section of clustered BP pairs similar to other threshold distances. Overall, because of the above biological reasons, we favored selecting a threshold distance that captures majority of the proximity-driven gene pair transcriptional correlations in these organisms as the basis for our BP-BP genome clustering analyses.

**Testing the statistical robustness of the threshold distance**

To confirm the statistical robustness of this threshold distance, we performed our clustering analyses (see below) and *p*-value quantifications for a range of different threshold distances (1X, 2X, 7X, 10X, 15X and 20X) and compared these *p*-values versus those generated using the 5X intergenic distance. This comparison was done for all organisms for all BP pairs. Figure S2 shows our analyses for the human genome as an example. In sum, these analyses revealed that significantly clustered BP pairs are robust at several different threshold distances in all model genomes analyzed in this report. Also, we compared the overlap of highly clustered BP pairs ($p<0.001$) among the different threshold values in each organism (Figure S3). These analyses revealed that 5X threshold distance captures a cross section of significantly clustered GO pairs in all organisms. Based on these data, we set 5X mean intergenic distance as the threshold distance for gene-pair clustering for the genomes used in this study.

**Determining the number of clustered gene pairs**

To quantify clustering between two BP terms (e.g., BP1 and BP2), we determined the number of instances genes from BP1 lie within the threshold distance of genes from BP2. Gene coordinates were used to count instances of gene pair clustering in BP1 and BP2.

To eliminate overestimation, if the same gene was found in the two BPs, the gene was removed from BP-BP clustering analysis.

**Specificity and performance of the statistical method used to quantify genome clustering**

To test the specificity of our statistical framework, we generated several random gene sets by selecting genes from the pool of BP-associated genes used in this study in each organism. To match the same gene number limits used to select BPs in each organism (see above), the number of randomly selected genes were 10-250 in yeast, 20-250 in worm and fly, and 50-250 in mouse and human. Next, all pairwise clustering analyses were performed and QQ plots were generated as depicted in Figure S4, revealing that randomly selected gene sets do not display significant genome clustering.

To monitor the performance of our statistical method, we selected 100 artificially generated gene sets and analyzed how the *p*-value for clustering changes as more clustered gene pairs were added incrementally to each analysis. As expected, the artificial addition of clustered gene pairs decreased the *p*-value for clustering (Figure S5). We also found that the magnitude of decrease in *p*-value caused by the incremental addition of clustered gene pairs to a given BP-BP analysis varies depending on the distribution of each gene set pair.

**Determining background clustering using Poisson distribution**

There are many previously developed methods for estimating genome clustering within a gene set or a functional group[22–24]; however, their utility for assessing clustering between BP pairs is limited. For example, they are not designed to eliminate intra-BP gene clustering (the tendency of genes in the same pathway to cluster) from the analysis. This necessitated the modification and development of a genome clustering platform such that the statistical significance of clustering between two BPs (e.g. BP1 and BP2) is quantified without contribution of intra-BP clustering. Background clustering was calculated for each BP-BP analysis in each organism. In total, over 2 million clustering analyses were performed in this study. To illustrate how background clustering between two BP terms was calculated, we use the following example: BP1 has 20 and BP2 has 30 genes. To determine background clustering between BP1 and BP2, first the 20 genes in BP1 were fixed in the genome and 30 genes were randomly selected 1,000 times from among the pool of genes with at least one ascribed BP designation. The numbers of clustered gene pairs generated from these 1,000 random samplings were then fit into a Poisson distribution. Next, the process was repeated by fixing the 30 genes in BP2, and 20 genes were picked randomly 1,000 times from among the pool of genes with at least one ascribed BP designation. The numbers of clustered gene pairs generated from these 1,000 random samplings were also fit into a Poisson distribution. In total, over 4 billion random samplings were performed to generate the Poisson graphs used in our study.

To calculate the significance of clustering for each pairwise analysis, we compared the observed number of clustered gene pairs versus the Poisson distribution generated by the background clustering as described above. Figure S6 shows simulated and fitted Poisson distributions for four representative sets of BP-BP analyses in the human genome. Note that the Poisson distribution can overestimate the *p*-value when the number of clustered gene pairs is large. In these instances, the fitted distribution is wider than the actual distribution (Figure S6D), thus this method is a stringent way to calculate *p*-values. This generates two *p*-values for each clustering analysis of which we used the smaller of the two. These values were used to generate the quantile-quantile (QQ) plots depicted in Figure 1D-H.

**Adjusting *p*-values and selecting highly clustered BP pairs**
Positive false discovery rate (pFDR)[25] was used to deal with the multiple comparison problem using a cutoff of 0.05. Also, we noted that some BP terms tended to cluster with many BP terms. To reduce this bias in our analyses, we kept only the top five most significantly clustered BP pairs in these instances. The list of significantly clustered BP pairs in all organisms can be found in Table S2. Significantly clustered BP pairs that had a Lin similarity score (see below for details) of 0.1 or less were considered 'disparate BP pairs'.

**Conservation of clustering**
To ask whether similar BP pairs cluster in the distantly related eukaryotes analyzed in this study, we selected BP terms which have clustered counterparts in at least two model organisms. The cutoff of two also permits the identification of mammalian-specific (mouse and human) or metazoan-specific (fly and worm) clustered BP pairs. This generated a list of 396 BP terms. Of the 396 BP terms 3 were absent in the R package and were not considered further. Using semantic similarity measurement scheme developed by Lin [26]and the R package GOSemSim[11], we found that the 393 BP terms form 30 highly similar BP groups (Table S3) (Figure 3A and Figure S7A). Highly similar BP groupings were defined by performing hierarchical clustering analysis on the Lin's similarity scores (ranging from 0-1.0) with the Euclidean pairwise distance (Figure 3A). The robustness of these BP groupings were tested against the Resnik semantic similarity method[27] revealing highly similar results (Figure S7A). We then asked in how many different genomes BPs in each group cluster with BPs in the other groups (Figure 3B and Figure S8).

We also used a parallel strategy in which we asked whether a given BP in Table S3 clusters with similar BP terms in the different genomes. For example, if BP1 clusters with BP2 and BP2' in mouse and yeast respectively, we used Lin to calculate the BP2- BP2' similarity score. The distribution of these values and the frequency with which highly similar (Lin scores >0.9) BP2-BP2' are found are depicted in Figure S7B.

**Human transcriptome analysis**
To quantify the transcriptional correlation between two BPs, we used the datasets available at the GTEx portal (v7 dataset) which contain transcriptomes of 53 different human tissues[5] from healthy individuals. The GTEx data were converted to mean $\log_2$ Transcript Per Million (TPM) values for each gene in every tissue (Figure S9A). Then the mean log2(TPM+1) expression for each gene was calculated across all 53 tissues. For each gene, these values were then normalized by subtracting the mean expression value in each tissue (Figure S9B). The normalized TPM values were used to calculate Z-scores

(deviation from the mean log TPM) for each gene across tissues (shown in Figure S9C). Using these data, we calculated transcriptional correlation between two BP terms as the correlation between mean tissue expression in BP1 and BP2 by summing all the Z-scores in each BP (53 mean tissue-specific Z-scores for each BP term as shown in Figure S9D).

**Calculating transcriptional correlations**

For transcriptional correlation calculations, we excluded genes that are present both in BP1 and BP2 from our calculations. Also, because clustered gene pairs are transcriptionally correlated (Figure S1), to remove this bias from our correlation analyses, we excluded the clustered gene pairs from all BP-BP correlation calculations. This permitted us to determine if the observed BP-BP correlations are driven by gene clusters or the entire pathway. Also, because similar GO terms often impact overlapping functions and are thus transcriptionally coordinated, to focus our analysis on disparate BPs, we separated BP pairs into two groups defined based on their semantic similarity score: (1) similar (Lin score >0.1) and (2) 'disparate BP pairs' (Lin score <=0.1)[26]. List of all BP pairs, the presence of shared clustered TFs, semantic similarity and transcriptional correlation scores can be found here (https://figshare.com/projects/Transcriptional_coherence/72644).

Figure 4 and Figure S10 illustrate the impact of clustered TF pairs on the transcriptional correlation of disparate (Lin score <=0.1) and similar (Lin score >0.1) pairs, respectively. 'circlize' package from the Comprehensive R Archive Network was used for draw the graph in Figure 4C[12]

**Human transcription factors**

The list of human TFs were obtained from UniProt [7].

**Clustering and transcriptional correlation analyses of biological pathway pairs defined by KEGG, BioCarta, and WikiPathways databases.**

To assess whether biological pathways defined by other databases also produce similar results to those found using Gene Ontology, we used pathway definitions provided by three different commonly used human databases, namely KEGG (N=186), BioCarta (N=292), and WikiPathways (N=615)[8].

Because there is no equivalent measure (semantic similarity) to quantify similarities between pathway pairs on these platforms, we defined 'disparate pathway pairs' as those which have no overlapping genes. To assess the effectiveness of this strategy in enriching for disparate BP pairs, we applied this strategy to the Gene Ontology BP pairs and found that BP pairs which share no overlapping genes have a lower Lin Score compared to those which share an overlapping gene (Figure S11A). In fact, 74% of nonoverlapping BP pairs have Lin similarity score of less than 0.1. Moreover, consistent with the observation that functionally similar BP pairs often impact overlapping functions and are thus transcriptionally coordinated, we found that BP pairs which share no overlapping genes have a lower transcriptional correlation compared to those which share at least one gene.

We followed the same analytical pipeline for identifying significantly clustered pathway pairs as those described in Figure 1B. The information about the pairwise analyses including the number of significantly clustered pathway pairs are summarized in Figure S12A. Also, the QQ plots for pairwise clustering within KEGG/BioCarta/Wiki-pathways (Figure S12B-D) were generated as described previously for GO (Figure 1D-H and Figure S4). In the end, the top 10% of the most significantly clustered pathway pairs based on their *p*-value were selected (KEGG (N=1,349), BioCarta (N=3,454), and WikiPathways (N=14,773), and separated into three bins bases on the presence of clustered TFs (Figure S12C-D). Again, because clustered gene pairs tend to be transcriptionally correlated (Figure S1), these genes were excluded from all transcriptional correlation calculations. This permitted us to identify functionally disparate pathway pairs whose transcriptional correlations are pathway-wide, and not driven by the coincidental correlation of their clustered gene pairs.

**Cancer cell line analysis**

Cancer Cell Line Encyclopedia (CCLE)[6] provide both gene expression and copy number variation data. For our analyses, we used 806 cell lines for which both transcriptome and copy number data were available. To estimate if a cancer cell line is an outlier in terms of the transcriptional coherence among cancer cell lines of the same type, we used the standardized residual of linear regression from a BP pair (BP1/BP2) within a specific tumor type. We calculated the mean expression of two BP terms for every cell line (excluding overlapping genes), and then used it to estimate two standardized residuals based on BP1 and BP2 as independent variables. The root mean squared values of two standardized residuals for TF-associated BP pairs (e.g. TF1 is associated with BP1, and TF2 is associated with BP2) and background (10,000 random BP pairs) are shown in Figure 5C-D and Figure S14C. This analysis was performed for cell lines that carry a deletion which eliminates one of the two clustered TFs or a translocation which physically separates a clustered TF pair.

**Topologically associated domains (TADs)**

To ask if the genes in a clustered TF pair occur in the same TAD, we used the 3D Genome Browser data and associated studies[17,28]. These datasets (http://promoter.bx.psu.edu/hi-c/publications.html) predict TADs by applying the Dixon et al.[17] pipeline to analyze their and other published datasets[29,30]. For each clustered TF pair, we calculated the number of times the clustered TFs were found in the same TAD in the 37 samples analyzed in this study. For Figure 5F and Figure S14, all the values sorted by x-values and y-values represent 50-points moving average.

**The maintenance of TF-TF clustering among 12 fly species**

We analyzed the genomes of 12 Drosophila species in FlyBase[10]. In the *D. melanogaster* genome, we defined clustered TF or gene pairs as those whose TF-TF or gene-gene distance fell within four times mean intergenic distance in the *D. melanogaster* genome. Next, we examined the intergenic distance of the corresponding gene or TF pair in the other fly species. Clustering was considered maintained if the gene-gene or TF-TF intergenic distance in the other Drosophila species also fell within four times mean intergenic distance in the other Drosophila genome. Figure 5G shows the number of fly species in which the gene-gene or TF-TF intergenic distance falls within the threshold distance (scale 0-11). The phylogenetic tree of fly species are shows in Figure S16, which was generated by TimeTree[13].

**Additional data**

Data for all pairwise analyses and MATLAB codes for this project can be found at https://figshare.com/projects/Transcriptional_coherence/72644.

### References


1. Wood, V., Harris, M.A., McDowall, M.D., Rutherford, K., Vaughan, B.W., Staines, D.M., Aslett, M., Lock, A., Bähler, J., Kersey, P.J., et al. (2012). PomBase: a comprehensive online resource for fission yeast. Nucl Acids Res *40*, D695-9. 10.1093/nar/gkr853.

2. Ashburner, M., Ball, C.A., Blake, J.A., Botstein, D., Butler, H., Cherry, J.M., Davis, A.P., Dolinski, K., Dwight, S.S., Eppig, J.T., et al. (2000). Gene ontology: tool for the unification of biology. Nat. Genet. *25*, 25–29. 10.1038/75556.

3. The Gene Ontology Consortium (2017). Expansion of the gene ontology knowledgebase and resources. Nucleic Acids Res. *45*, D331–D338. 10.1093/nar/gkw1108.

4. Zerbino, D.R., Achuthan, P., Akanni, W., Amode, M.R., Barrell, D., Bhai, J., Billis, K., Cummins, C., Gall, A., Girón, C.G., et al. (2018). Ensembl 2018. Nucleic Acids Res. *46*, D754–D761. 10.1093/nar/gkx1098.

5. The GTEx Consortium (2013). The Genotype-Tissue Expression (GTEx) project. Nat. Genet. *45*, 580–585. 10.1038/ng.2653.

6. Barretina, J., Caponigro, G., Stransky, N., Venkatesan, K., Margolin, A.A., Kim, S., Wilson, C.J., Lehár, J., Kryukov, G. V., Sonkin, D., et al. (2012). The Cancer Cell Line Encyclopedia enables predictive modelling of anticancer drug sensitivity. Nature *483*, 603–607. 10.1038/nature11003.

7. Bateman, A., Martin, M.J., O'Donovan, C., Magrane, M., Alpi, E., Antunes, R., Bely, B., Bingley, M., Bonilla, C., Britto, R., et al. (2017). UniProt: the universal protein knowledgebase. Nucleic Acids Res. *45*, D158–D169. 10.1093/nar/gkw1099.

8. Liberzon, A., Subramanian, A., Pinchback, R., Thorvaldsdóttir, H., Tamayo, P., and Mesirov, J.P. (2011). Molecular signatures database (MSigDB) 3.0. Bioinformatics *27*, 1739–1740. 10.1093/bioinformatics/btr260.

9. Subramanian, A., Tamayo, P., Mootha, V.K., Mukherjee, S., Ebert, B.L., Gillette, M.A., Paulovich, A., Pomeroy, S.L., Golub, T.R., Lander, E.S., et al. (2005). Gene set enrichment analysis: A knowledge-based approach for interpreting genome-wide

expression profiles. Proc. Natl. Acad. Sci. *102*, 15545–15550. 10.1073/pnas.0506580102.

10. Larkin, A., Marygold, S.J., Antonazzo, G., Attrill, H., dos Santos, G., Garapati, P.V., Goodman, J.L., Gramates, L.S., Millburn, G., Strelets, V.B., et al. (2021). FlyBase: updates to the Drosophila melanogaster knowledge base. Nucleic Acids Res. *49*, D899–D907. 10.1093/nar/gkaa1026.

11. Yu, G., Li, F., Qin, Y., Bo, X., Wu, Y., and Wang, S. (2010). GOSemSim: an R package for measuring semantic similarity among GO terms and gene products. Bioinformatics *26*, 976–978. 10.1093/bioinformatics/btq064.

12. Gu, Z., Gu, L., Eils, R., Schlesner, M., and Brors, B. (2014). circlize implements and enhances circular visualization in R. Bioinformatics *30*, 2811–2812. 10.1093/bioinformatics/btu393.

13. Kumar, S., Stecher, G., Suleski, M., and Hedges, S.B. (2017). TimeTree: A Resource for Timelines, Timetrees, and Divergence Times. Mol. Biol. Evol. *34*, 1812–1819. 10.1093/molbev/msx116.

14. Joh, R.I., Khanduja, J.S., Calvo, I.A., Mistry, M., Palmieri, C.M.C.M., Savol, A.J.A.J., Ho Sui, S.J.S.J., Sadreyev, R.I.R.I., Aryee, M.J.M.J., and Motamedi, M. (2016). Survival in quiescence requires the euchromatic deployment of Clr4/SUV39H by Argonaute-associated small RNAs. Mol. Cell *64*, 1088–1101. 10.1016/j.molcel.2016.11.020.

15. Thévenin, A., Ein-Dor, L., Ozery-Flato, M., and Shamir, R. (2014). Functional gene groups are concentrated within chromosomes, among chromosomes and in the nuclear space of the human genome. Nucleic Acids Res. *42*, 9854–9861. 10.1093/nar/gku667.

16. Mizuguchi, T., Fudenberg, G., Mehta, S., Belton, J.-M., Taneja, N., Folco, H.D., FitzGerald, P., Dekker, J., Mirny, L., Barrowman, J., et al. (2014). Cohesin-dependent globules and heterochromatin shape 3D genome architecture in S. pombe. Nature *516*, 432–435. 10.1038/nature13833.

17. Dixon, J.R., Selvaraj, S., Yue, F., Kim, A., Li, Y., Shen, Y., Hu, M., Liu, J.S., and Ren, B. (2012). Topological domains in mammalian genomes identified by analysis of chromatin interactions. Nature *485*, 376–380. 10.1038/nature11082.

18. Weber, C.C., and Hurst, L.D. (2011). Support for multiple classes of local expression clusters in Drosophila melanogaster, but no evidence for gene order conservation. Genome Biol. *12*, R23. 10.1186/gb-2011-12-3-r23.

19. Spellman, P.T., and Rubin, G.M. (2002). Evidence for large domains of similarly expressed genes in the Drosophila genome. J. Biol. *1*, 5.

20. Szabo, Q., Bantignies, F., and Cavalli, G. (2019). Principles of genome folding into topologically associating domains. Sci. Adv. *5*, eaaw1668. 10.1126/sciadv.aaw1668.

21. Ramírez, F., Bhardwaj, V., Arrigoni, L., Lam, K.C., Grüning, B.A., Villaveces, J., Habermann, B., Akhtar, A., and Manke, T. (2018). High-resolution TADs reveal DNA sequences underlying genome organization in flies. Nat. Commun. *9*, 189. 10.1038/s41467-017-02525-w.

22. Lercher, M.J., Urrutia, A.O., and Hurst, L.D. (2002). Clustering of housekeeping genes provides a unified model of gene order in the human genome. Nat. Genet. *31*, 180–183. 10.1038/ng887.

23. Versteeg, R., van Schaik, B.D.C., van Batenburg, M.F., Roos, M., Monajemi, R., Caron, H., Bussemaker, H.J., and van Kampen, A.H.C. (2003). The human transcriptome map reveals extremes in gene density, intron length, GC content, and repeat pattern for domains of highly and weakly expressed genes. Genome Res. *13*, 1998–2004. 10.1101/gr.1649303.

24. Caron, H., van Schaik, B., van der Mee, M., Baas, F., Riggins, G., van Sluis, P., Hermus, M.C., van Asperen, R., Boon, K., Voûte, P.A., et al. (2001). The human transcriptome map: clustering of highly expressed genes in chromosomal domains. Science *291*, 1289–1292. 10.1126/science.1056794.

25. Storey, J.D. (2003). The positive false discovery rate: a Bayesian interpretation and the q -value. Ann. Stat. *31*, 2013–2035. 10.1214/aos/1074290335.

26. Lin, D. (1998). An information-theoretic definition of similarity. Proc. Fifteenth Int. Conf. Mach. Learn., 296–304.

27. Resnik, P. (1995). Using information content to evaluate semantic similarity in a taxonomy. Proc. 14TH Int. Jt. Conf. Artif. Intell. *1*, 448--453.

28. Wang, Y., Song, F., Zhang, B., Zhang, L., Xu, J., Kuang, D., Li, D., Choudhary, M.N.K., Li, Y., Hu, M., et al. (2018). The 3D genome browser: a web-based browser for visualizing 3D genome organization and long-range chromatin interactions. Genome Biol. *19*, 151. 10.1186/s13059-018-1519-9.

29. Lieberman-Aiden, E., van Berkum, N.L., Williams, L., Imakaev, M., Ragoczy, T., Telling, A., Amit, I., Lajoie, B.R., Sabo, P.J., Dorschner, M.O., et al. (2009). Comprehensive mapping of long-range interactions reveals folding principles of the human genome. Science *326*, 289–293. 10.1126/science.1181369.

30. Leung, D., Jung, I., Rajagopal, N., Schmitt, A., Selvaraj, S., Lee, A.Y., Yen, C.A., Lin, S., Lin, Y., Qiu, Y., et al. (2015). Integrative analysis of haplotype-resolved epigenomes across human tissues. Nature *518*, 350–354. 10.1038/nature14217.
