## Supplementary figures for "Gene clustering drives the transcriptional coherence of disparate biological processes in eukaryotes"

### Supplemental Figures

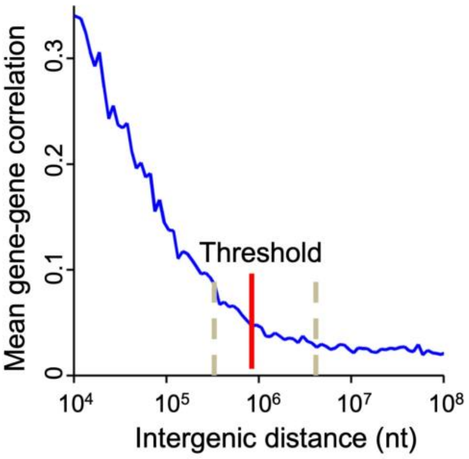


Figure S1

**Figure S1.**

**Transcriptional correlation of human gene pairs is a function of their intergenic distance.** The graph depicts the transcriptional correlation of all possible human gene pairs as a function of the pairwise intergenic distance using the human GTEx data. 200 intergenic distance bins were created ranging from 20-230MB and the mean transcriptional correlation of all gene pairs within each bin was calculated and plotted (blue line). The red line represents the threshold distance (five times (5X) mean intergenic distance) chosen to define clustered gene pairs in the human genome in this paper (see Materials and Methods). Left and right dashed gray lines represent 2X and 20X mean intergenic distances, respectively, shown for comparison.

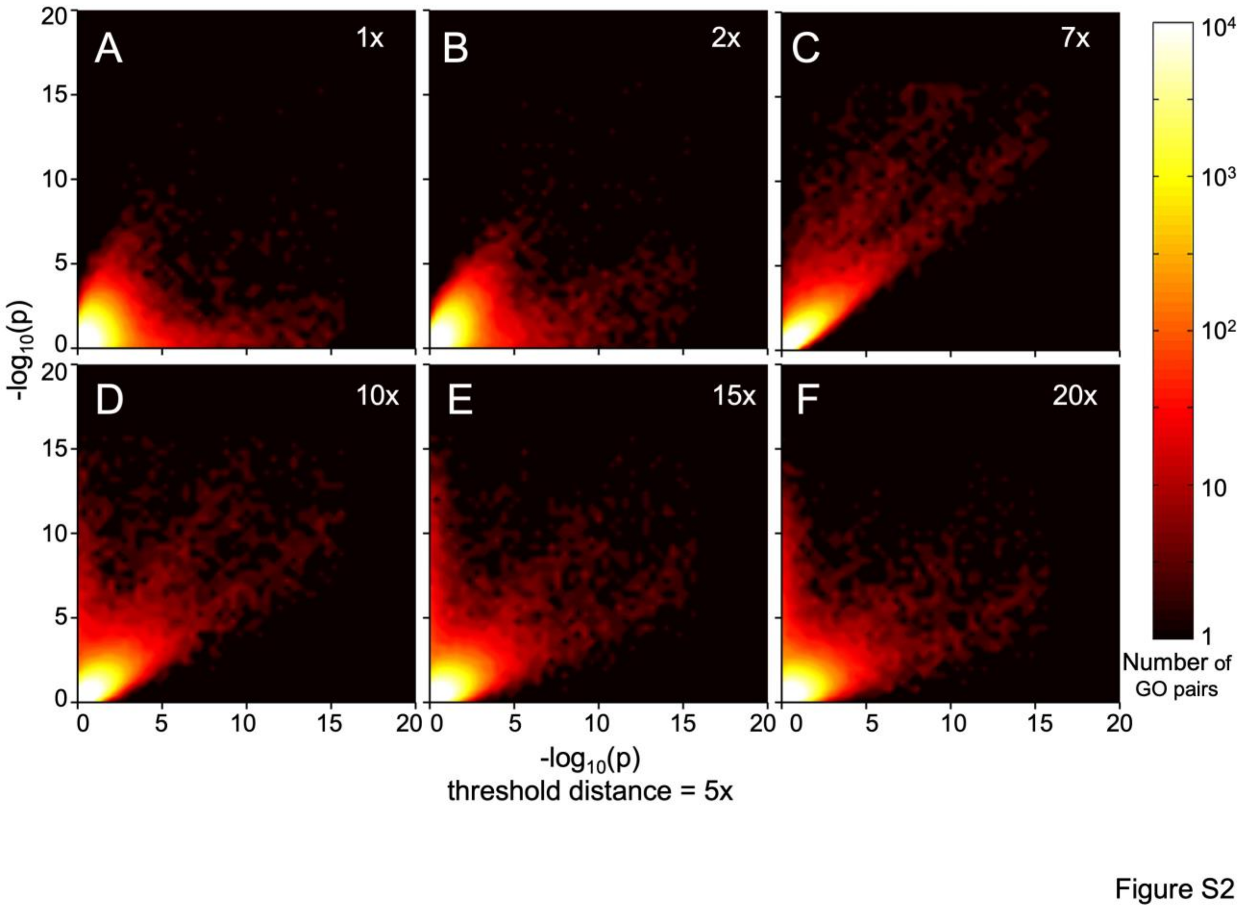


**Figure S2.**

**5X mean intergenic distance is robust for identifying clustered BP pairs in the human genome.** Heatmaps depicting distribution of *p*-values ($\log_{10}(p)$) for identifying significantly clustered BP pairs calculated for five times (5X) mean intergenic distance versus the *p*-values calculated for the indicated threshold distances (1X, 2X, 7X, 10X, 15X, 20X mean intergenic distance). Color represent the significance of clustering between BP pairs. The data for other model organisms are available at https://figshare.com/projects/Transcriptional_coherence/72644.

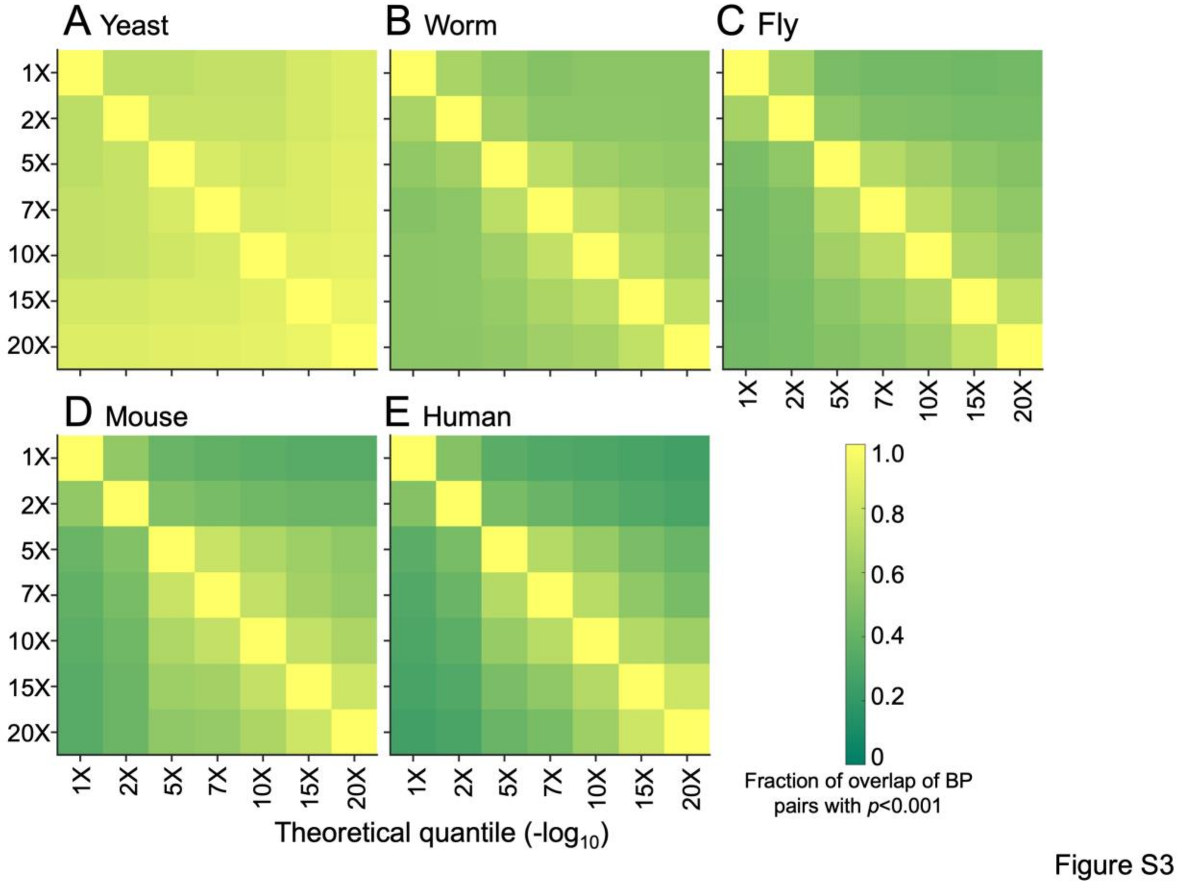


**Figure S3.**
**5X threshold distance captures a cross section of significantly clustered BP pairs in all organisms.** Heat map depicting the fraction of overlap of significantly clustered BP pairs with *p* values less than 0.001 for the indicated threshold distances (1X, 2X, 5X, 7X, 10X, 15X, 20X mean intergenic distance) in each organism. Color is proportional to the fraction of overlap of significantly clustered BP pairs at each threshold distance.

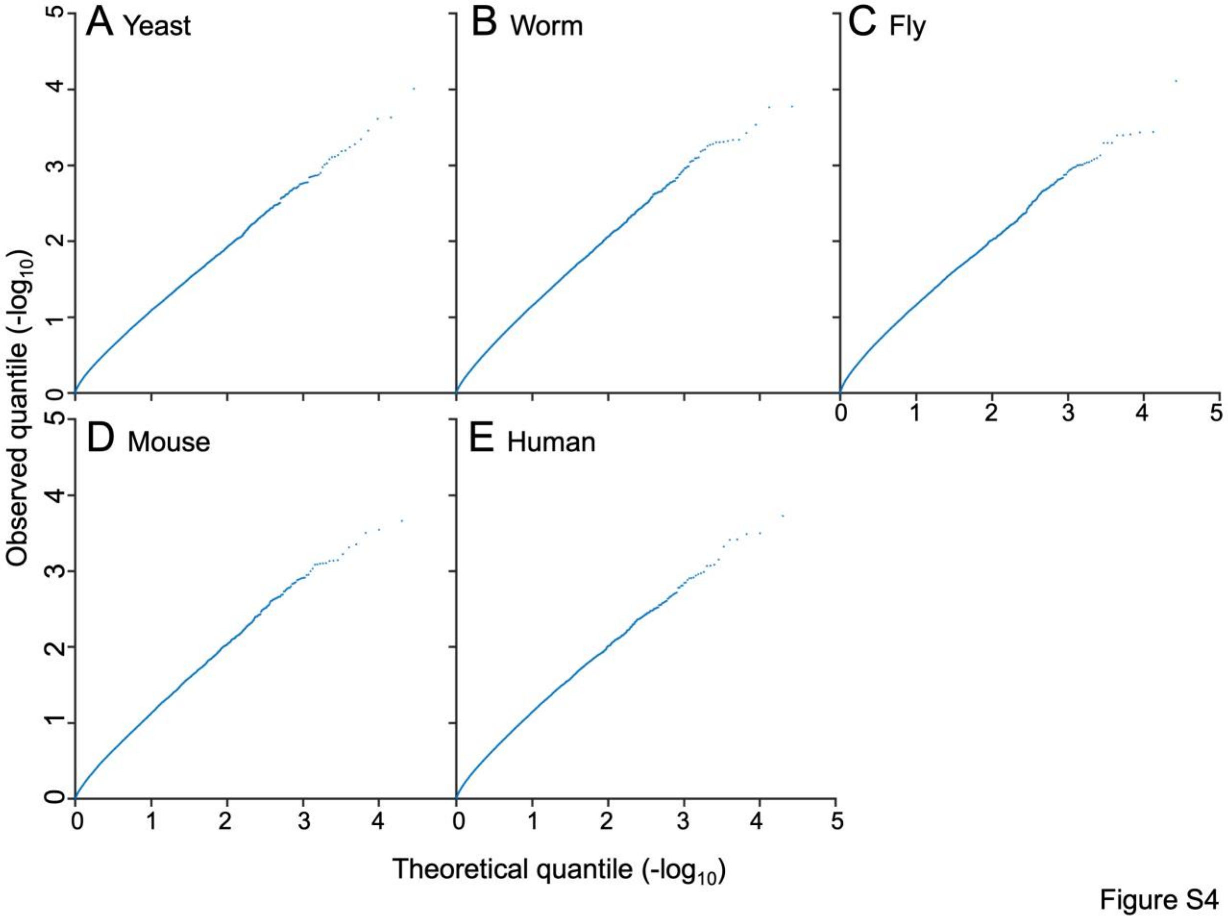


**Figure S4.**

**Randomly selected gene sets do not display significant genome clustering.** Quantile-quantile (QQ) plots of observed (Y-axis) versus theoretical (X-axis) *p*-value distributions for genome clustering between gene sets generated by randomly selecting genes from the pool of BP-associated genes in each organism (see Materials and Methods). The QQ plots in (A) yeast (*S. pombe*), (B) worm (*C. elegans*), (C) fly (*D. melanogaster*), (D) mouse (*M. musculus*) and (E) human (*H. sapiens*) are depicted. In contrast to naturally occurring BPs (Figure 1D to H), these QQ plots do not curl up, thus fail to exhibit significant clustering. These analyses reveal that our statistical method specifically identifies significantly clustered BP pairs in each genome.

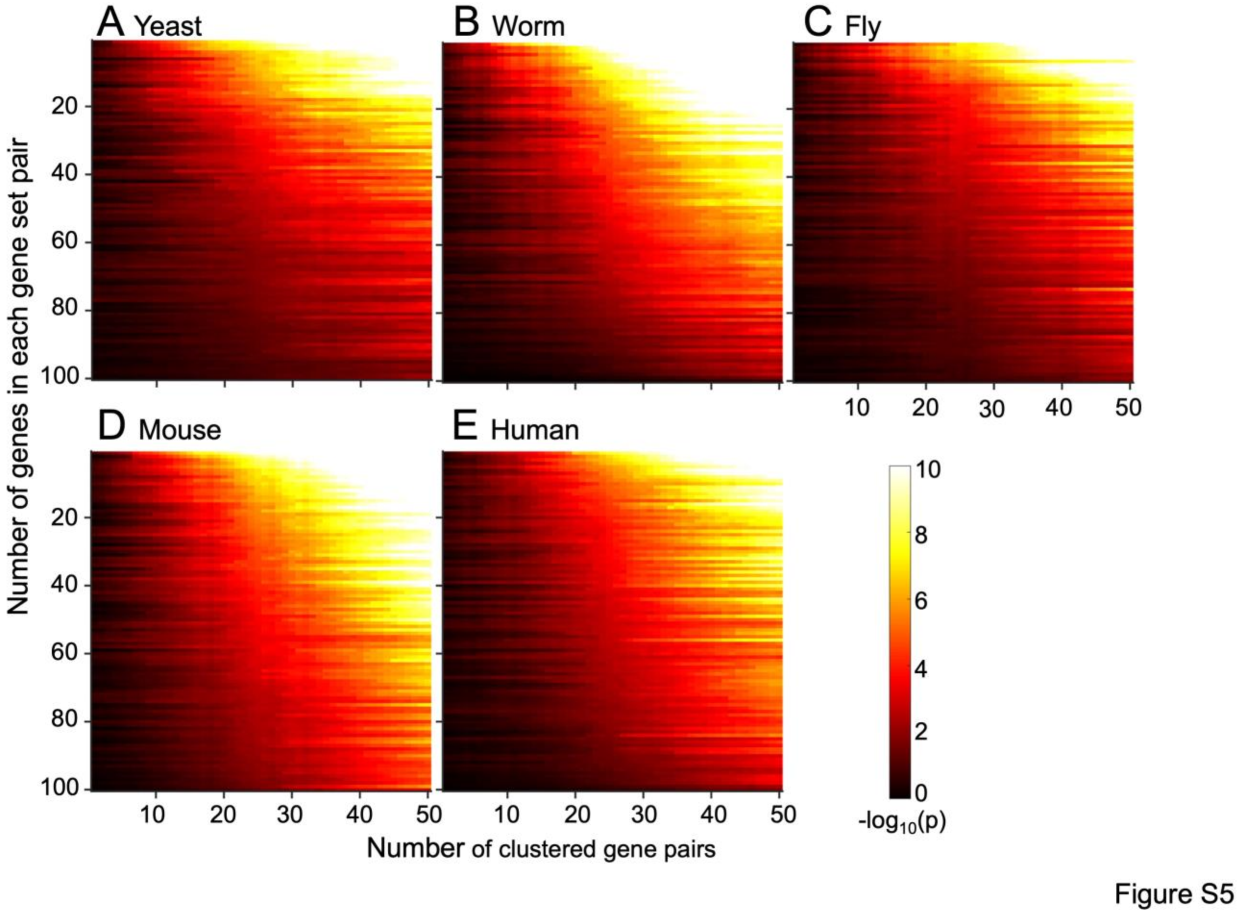


**Figure S5.**

**The statistical method developed for genome clustering performs as expected.** Heat maps depicting the impact on *p*-value as clustered gene pairs are artificially added to randomly generated gene sets. For each indicated organism, *p*-values for clustering of 100 artificially generated pairs of gene sets were calculated. As expected, the artificial addition of clustered gene pairs decreases the *p*-value for clustering in all model organisms. Color is proportional to $-\log_{10}(p)$.

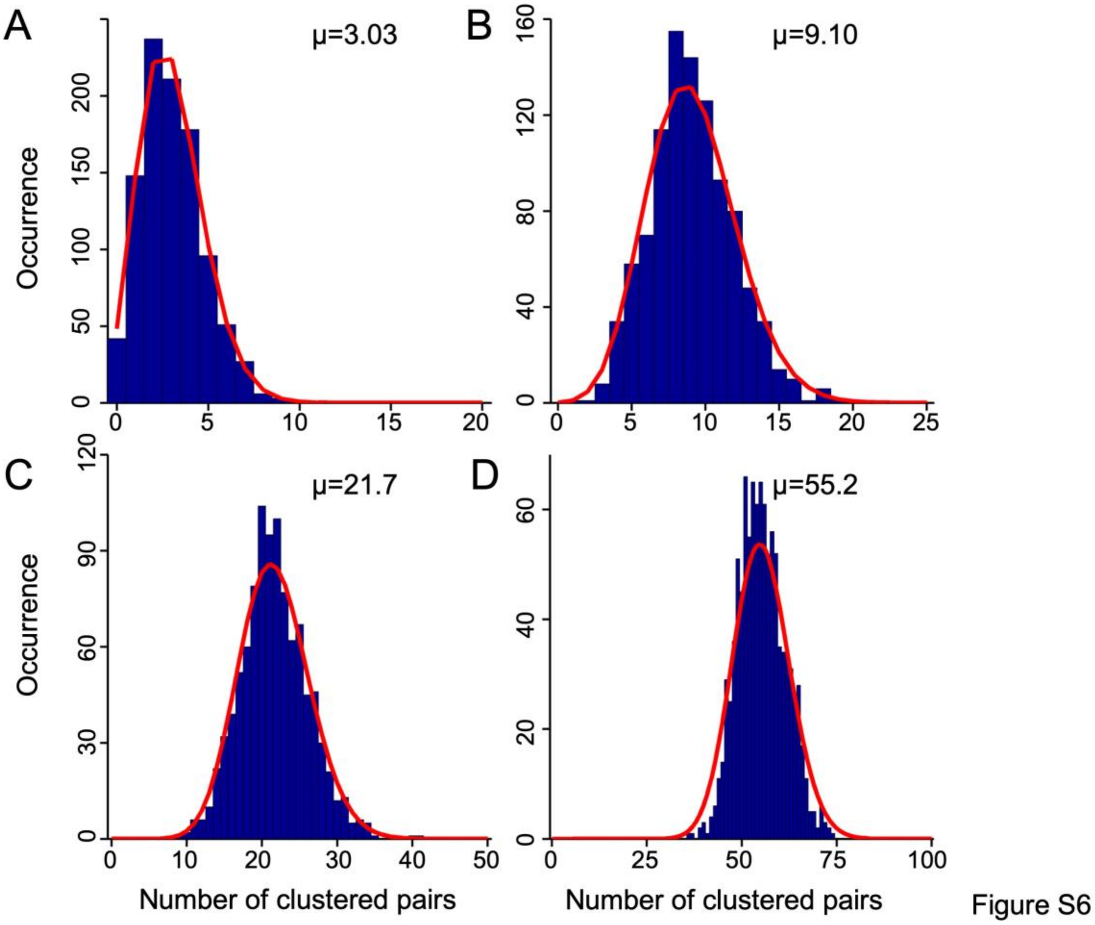


**Figure S6.**

**Examples of clustering quantified by random gene sampling.** Graphs depicting the distribution of background gene clustering occurrences (bar graphs) and their Poisson fit (red lines) for four representative sets of BP-BP analyses performed on the human genome. 2,000 random samplings (StarMethods) were done to quantify background gene clustering for each BP pair analysis in each organism. μ is the mean number of clustered gene pairs derived from the fitted Poisson distribution.

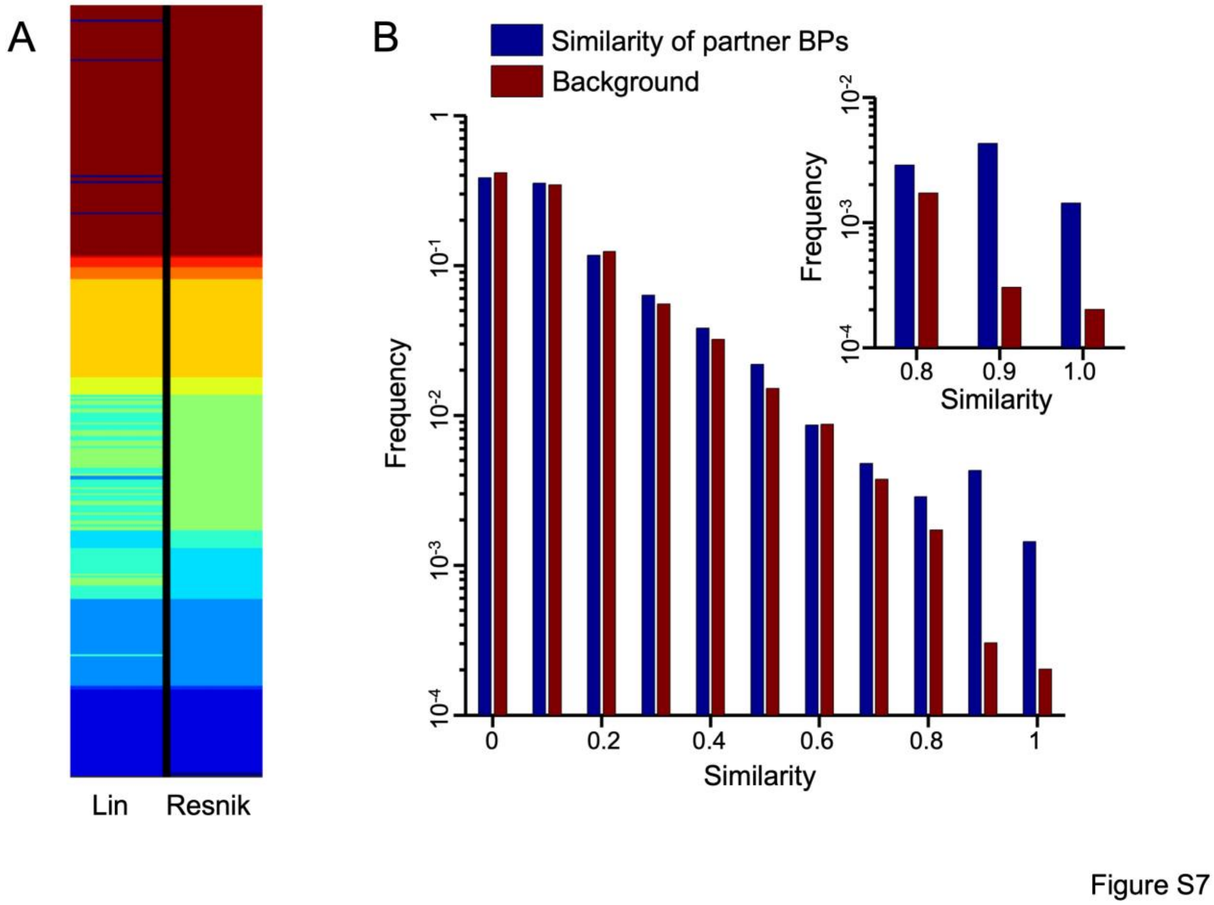


**Figure S7.**

**Highly similar BP pairs are clustered in eukaryotes**. (A) Graph depicting the BP groups generated by hierarchical clustering using Lin's and Resnik's semantic similarity methods. The BPs used in this analysis all have a clustered BP partner in at least two model organisms. (B) Graph depicting the frequency of semantic similarity score of BP terms which share a common clustered partner in at least two organisms. For example if BP1 clusters with BP2 and BP2' in mouse and yeast genomes, respectively, here we calculated the similarity bewteen BP2 and BP2' using the Lin method.

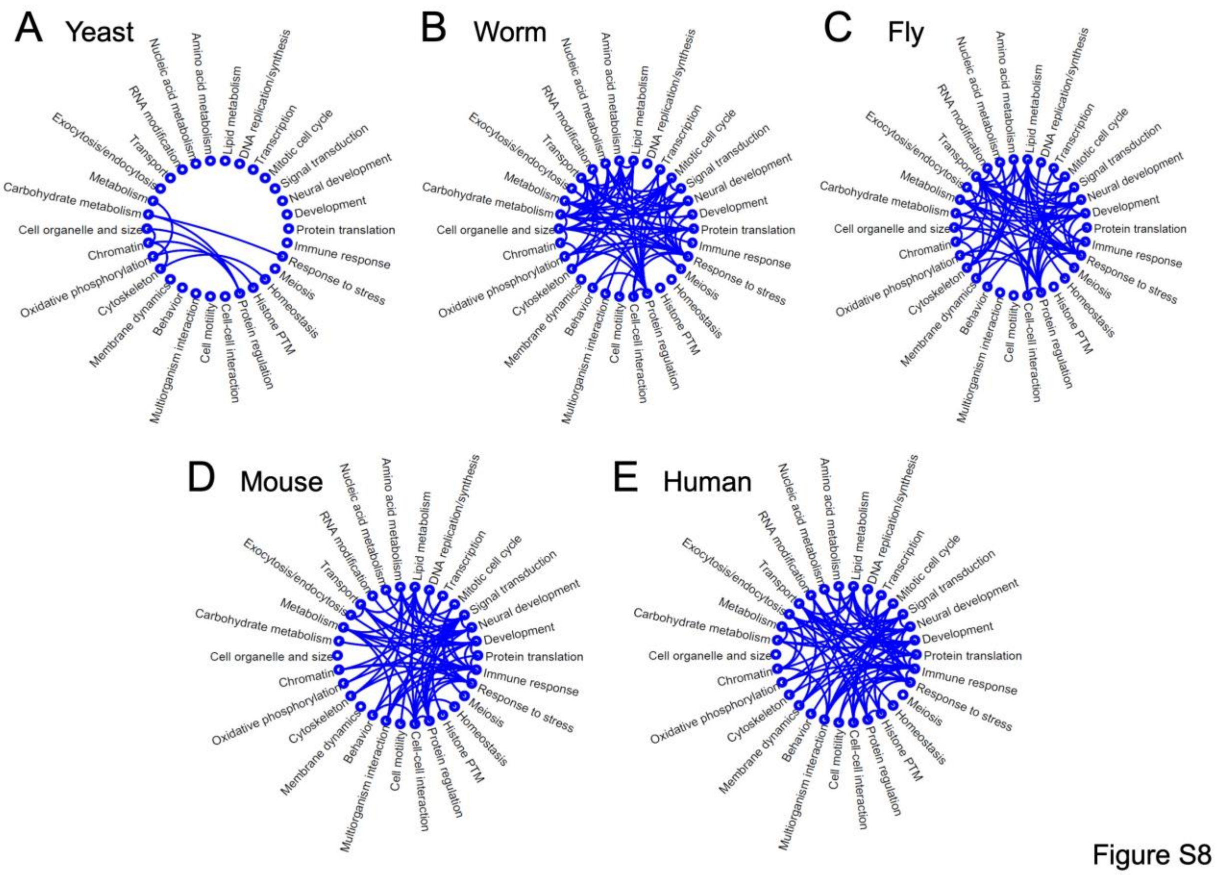


**Figure S8.**
**Several BP groups cluster in multiple eukaryotic genomes.** The lines connect two BP groups which have at least one clustered BP pair between them in the indicated organisms.

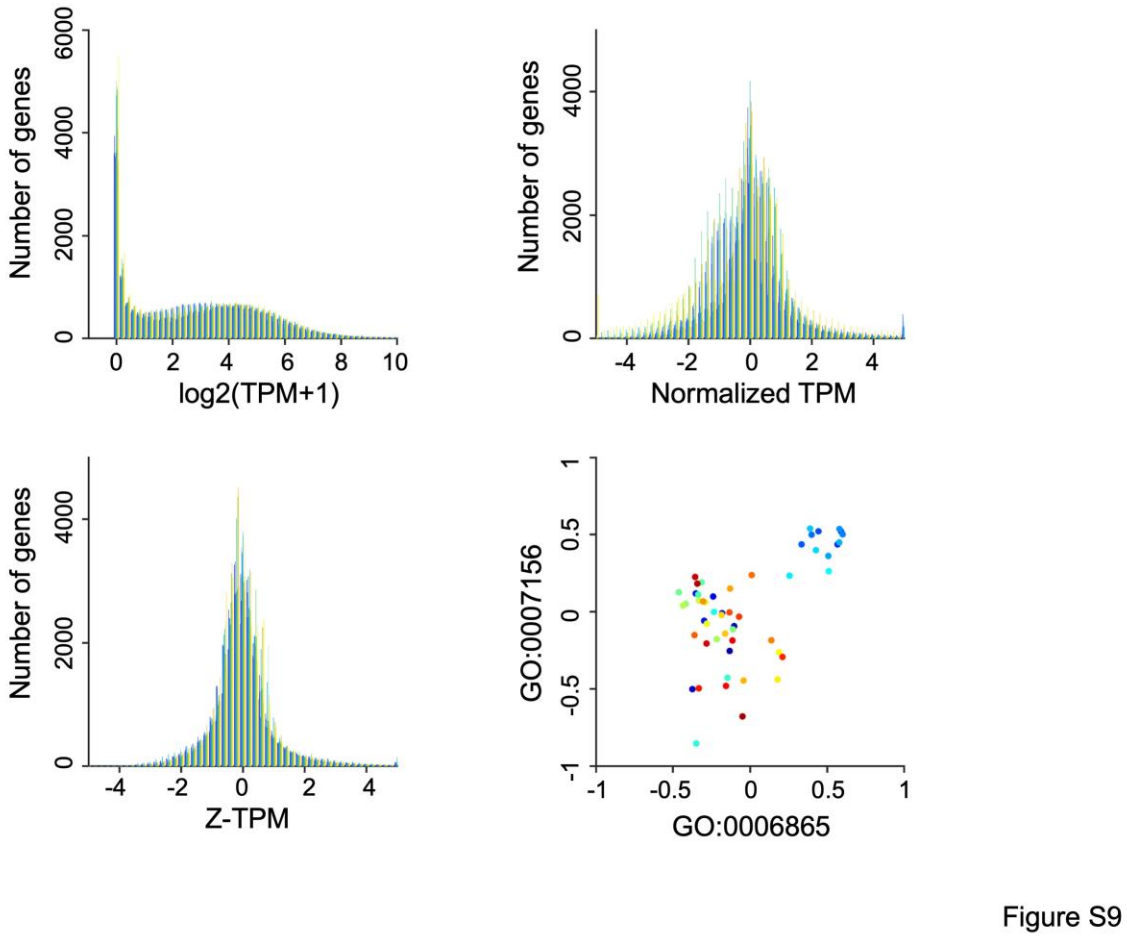


**Figure S9.**
**Pre-processing of GTEx transcriptome data to quantify transcriptional correlation between BP pairs.** Graphs depict the (A) raw Transcript Per Million (TPM) data ($\log_2$(TPM+1)), (B) normalized TPM and (C) Z-score of TPM. Each color corresponds to a different tissue. The Z-scores were used for quantitative comparisons across different tissues. (D) An example of BP-BP transcriptional correlation in human (GO:0006959 (humoral Immune response) and GO:0007162 (negative regulation of cell adhesion)). Each dot represents a different tissue in GTEx.

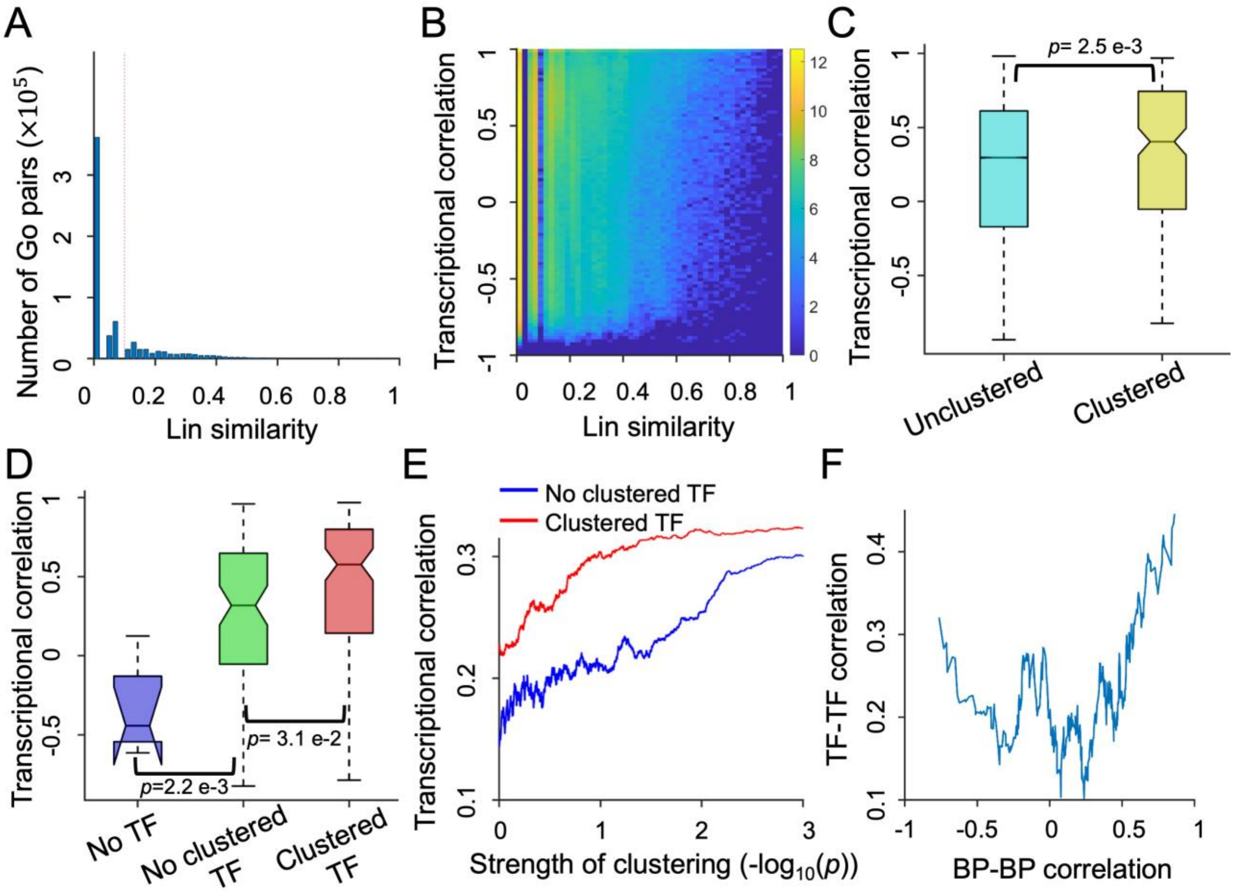


**Figure S10.**

**Functionally similar (fs) clustered BP pairs (Lin score > 0.1), especially those with clustered TFs, are transcriptionally correlated in human cells.** (A) Graph depicting the distribution of Lin similarity score among all human BP pairs analyzed in this study. The dotted red line represents Lin Score of 0.1. (B) Heat map depicting the distribution of Lin similarity score and transcriptional correlation of the associated BP pairs. The color is proportional to the number of BP pairs (log2). (C) Box plots depicting transcriptional correlation of clustered (N=188) and unclustered (N=186,330) fs-BP pairs. (D) Box plots depicting transcriptional correlation of clustered fs-BP pairs assigned to three groups based on the presence and clustering of TFs. 'No TF' refers to clustered fs-BP pairs in which one or both of the two BP terms is missing a TF (N=7); 'No clustered TF' refers to clustered fs-BP pairs in which both BP terms have a TF, but their TFs are not clustered (N=78); and 'Clustered TF' refers to clustered fs-BP pairs in which both BP pairs have a TF and those TFs are clustered (N=103). *p*-values were calculated by the two-sample t-test. (E) Graph depicting the transcriptional correlation of all human fs-BP pairs as a function of the strength of their genomic clustering. Red and blue lines represent transcriptional correlation of BP pairs with (N=27,898) or without (N=158,016) clustered TFs, respectively. Each BP pair was sorted by the strength of clustering and the correlation represents the moving average of 5,000 points. (F) Graph depicting TF-TF transcriptional correlation as a function of the transcriptional correlation of the associated fd-BP pairs. TF-TF pairs were sorted by BP-BP correlation, and the moving averages of 50 points are shown.

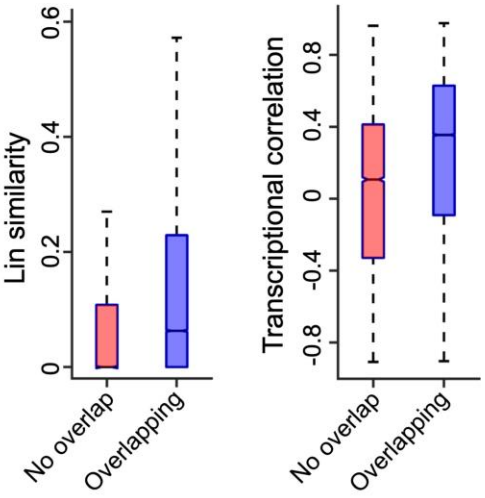


Figure S11

**Figure S11.**

**BP pairs sharing no overlapping genes display lower Lin score and transcriptional correlation compared to BP pairs which share at least one gene.** (A) Box plots depicting the Lin similarity of BP pairs which share (overlapping) or do not share (no overlap) at least one gene. 74% of ‘no overlap’ BP pairs have Lin similarity score of less than 0.1. (B) Transcriptional correlation of BP pairs which share (overlapping) or do not share (no overlap) at least one gene. BPs were defined by Gene Ontology in these analyses.

A

| | KEGG | BioCarta | Wiki pathways |
|---|---|---|---|
| Number of pathways | 186 | 292 | 615 |
| Number of pairwise analyses | 17,205 | 42,486 | 188,805 |
| Number of nonoverlapping pathway pair analyses | 13,626 | 34,147 | 140,277 |
| Number of clustered pathway pairs p<0.05 | 2,221 | 7,207 | 23,337 |
| Number of clustered pathway pairs q<0.05 | 546 | 538 | 771 |

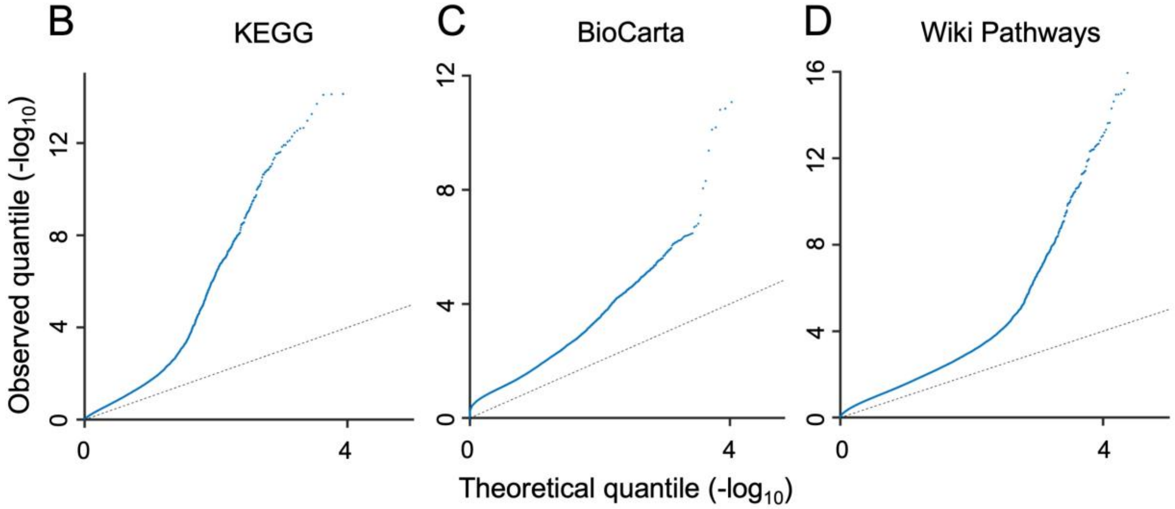


Figure S12

**Figure S12.**

**Application of BP-BP clustering pipeline to biological pathways defined by platforms other than Gene Ontology reveal clustering of several hundred disparate pathway pairs in the human genome.** (A) Table summarizing the number of pathway pairs, pairwise analyses, non-overlapping pathway pairs, and significantly clustered pathway pairs identified in the human genome. The analyses were performed on biological pathways as defined by KEGG, BioCarta and Wiki Pathways. (B-D) Quantile-quantile (QQ) plots of observed (Y-axis) versus theoretical (X-axis) *p*-value distributions for clustering between pathways pairs (see StarMethods). The QQ plots in (B) KEGG , (C) BioCarta, and (D) Wiki pathways are depicted.

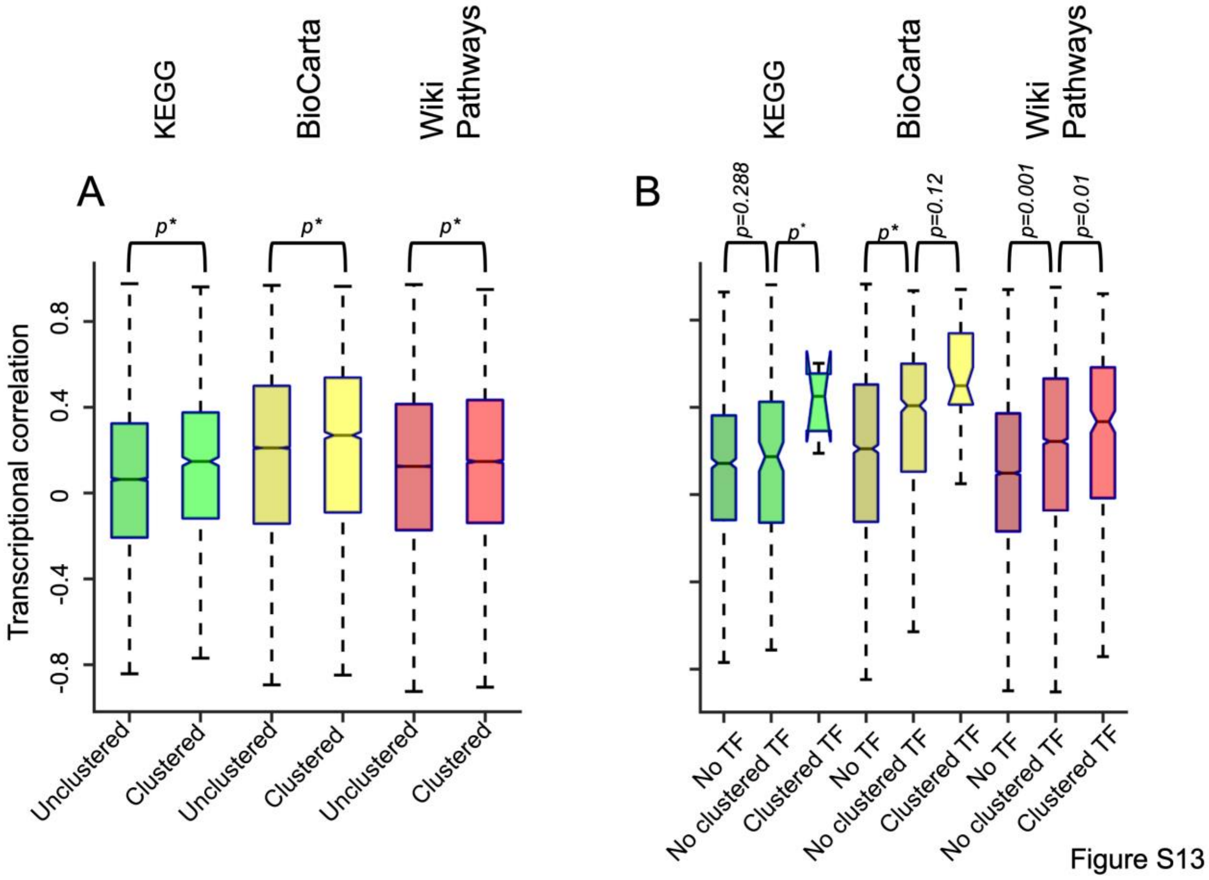


**Figure S13.**

**Clustered pathway pairs, especially those with a clustered TF, are transcriptionally correlated in human.** (A) Box plots depicting transcriptional correlation of clustered and unclustered pathway pairs. The analyses were performed on disparate pathway pairs which we defined as those which share no overlapping genes (see Figure S11). Clustering was quantified as described in StarMethods. (B) Box plots depicting transcriptional correlation of clustered pathway pairs assigned to three groups based on the presence and clustering of TFs. ‘No TF’ refers to clustered pathway pairs in which one or both pathways are missing a TF; ‘No clustered TF’ refers to clustered pathway pairs in which both pathways have a TF, but their TFs are not clustered; and ‘Clustered TF’ refers to clustered pathway pairs in which both pathway pairs have a TF and share at least one clustered TF pair. *p*-values were calculated by the two-sample t-test.

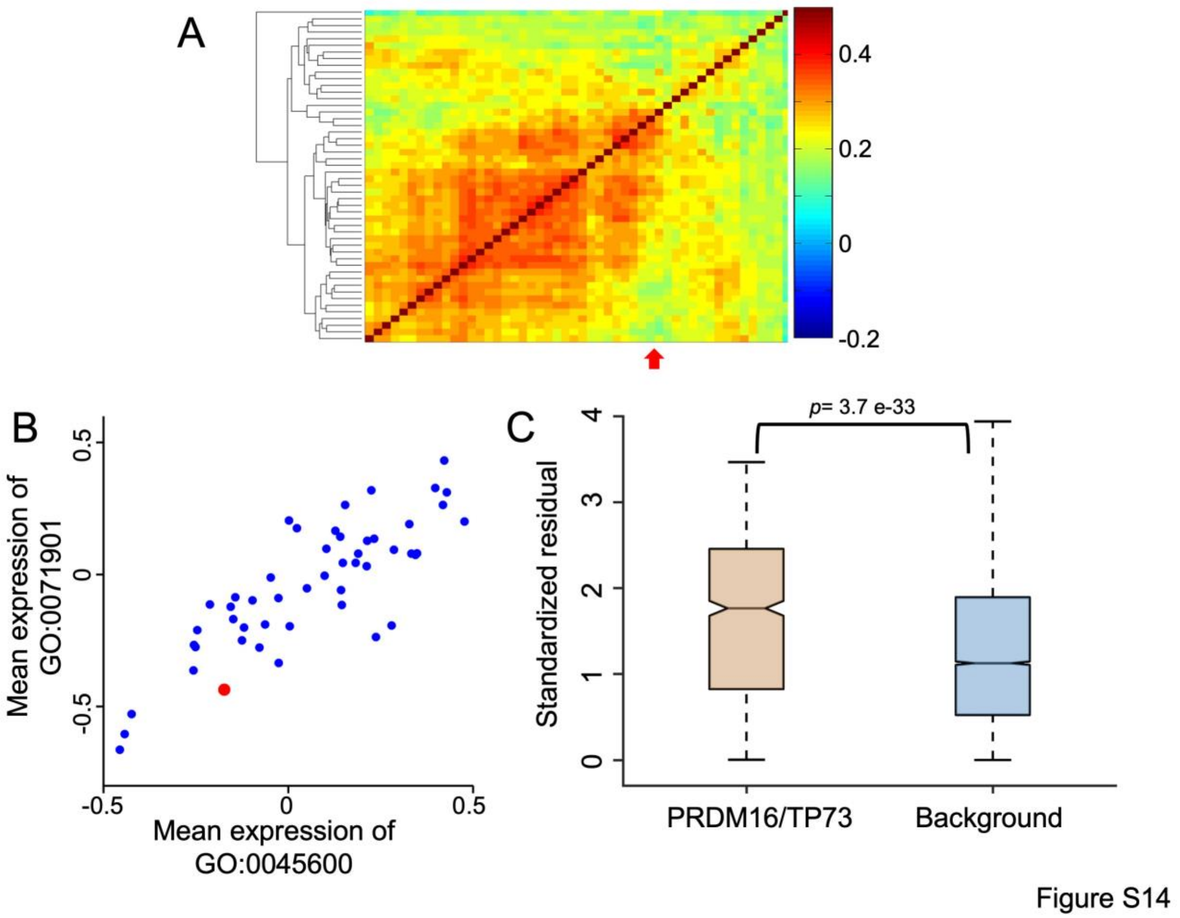


**Figure S14.**
**Transcription factor (TF) clustering drives the transcriptional coregulation of fd-BP pairs.**
(A) Hierarchical clustering of transcriptomes of transverse colon cancer cell lines (N=50). Red arrow indicates SNU1033, the cell line carrying a deletion in PRDM16. The color indicates to the overall pairwise correlation between two cancer cell lines. (B) Graph depicting the mean expression of two PRDM16- and TP73-associated BPs (GO:0045600 positive regulation of fat cell differentiation and GO:0071901 negative regulation of protein serine/threonine kinase activity, respectively) among transverse colon cancer cell lines. Red dot represents SNU1033. (C) Boxplot depicting the distribution of standardized residuals of BP pairs in SNU1033 versus other transverse colon cancer cell lines. The left and right bars depict this distribution for the PRDM16- and TP73-associated BP pairs and random BP pairs, respectively.

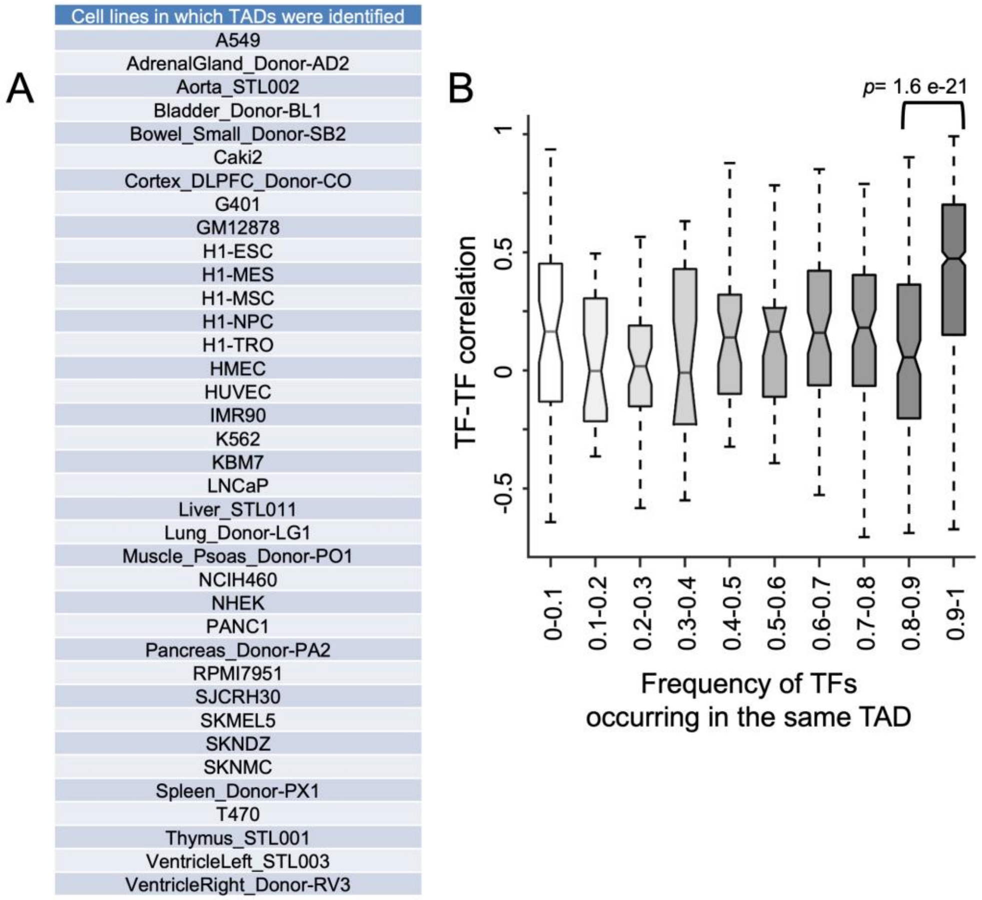


Figure S15

**Figure S15.**

**Clustered TFs found frequently in the same TAD display high TF-TF transcriptional correlation.** (A) The list of 37 cell lines in which TADs were identified. This was used to calculate the frequency by which a clustered TF pairs falls in the same TAD based on these studies. (B) Graph depicting TF-TF correlation and the frequency which clustered TFs are found in the same TAD. *p*-values were calculated by the two-sample t-test.

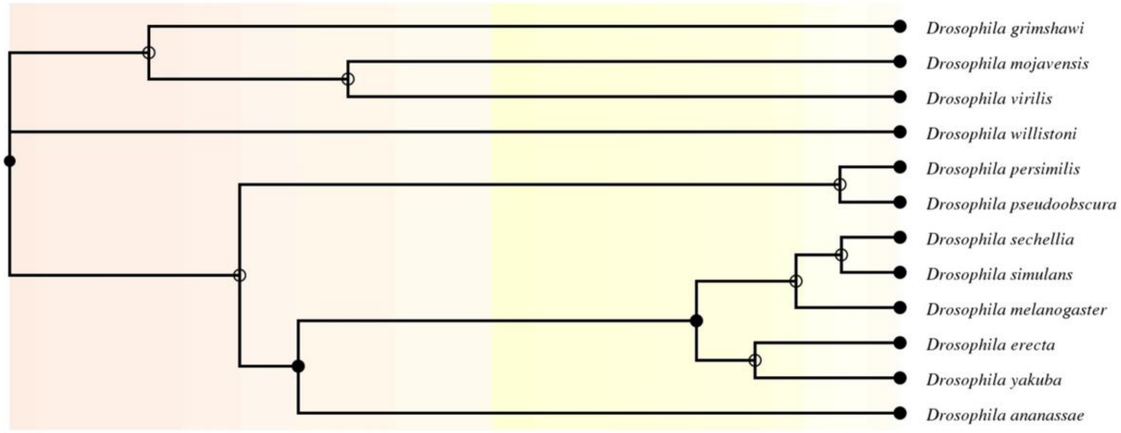


Figure S16

**Figure S16.**
**Phylogenic evolutionary relationship of 12 species among the Drosophila genus by TimeTree.** See StarMethods for details.
